# The ganglioside profile of mouse brain does not respond to changes in dietary n-3 polyunsaturated fatty acid supply

**DOI:** 10.64898/2026.11.06.762811

**Authors:** E.A.Y. Masson, L. Martine, S. Cabaret, S. Gregoire, F. Ducrocq, P. Trifilieff, N. Acar, O. Berdeaux

## Abstract

Long chain polyunsaturated fatty acids (LC-PUFAs) of the n-3 and n-6 families are major components of brain lipids, both in terms of levels and biological significance. They play crucial roles in various neurological processes. The brain is largely dependent on food supply of LC-PUFAs and its lipid composition can be modulated by changes in dietary intake. Sufficient dietary supply of the n-3 LC-PUFAs as well as a balanced n-6/n-3 ratio are required for proper brain development and function. As dietary intervention using n-3 LC-PUFAs is considered a promising therapeutic approach to prevent or treat brain disorders, it is important to understand the associated biochemical changes as well as the mechanisms underlying the biological effects. In the present study, we performed lipidomic analyses to document the effects of dietary n-3 LC-PUFA supply on the composition of two families of FA-carrying complex membrane lipids playing important roles in the nervous system: phospholipids and gangliosides. We show that an unbalanced, n-3 LC-PUFA-deficient diet, from gestation to adulthood in mice resulted in decreased levels of docosahexaenoic acid (DHA, C22:6n-3) and increased levels of docosapentaenoic acid (DPA, C22:5n-6) in brain phospholipids, while a n-3 LC-PUFA-supplemented diet was associated with the reverse phenotype. However, the composition of brain gangliosides, which carry mainly saturated and mono-unsaturated fatty acids, was unaffected by the diet, suggesting that they do not directly contribute to mediating the neurological effects of dietary n-3 LC-PUFA manipulation.

**Graphical abstract:** 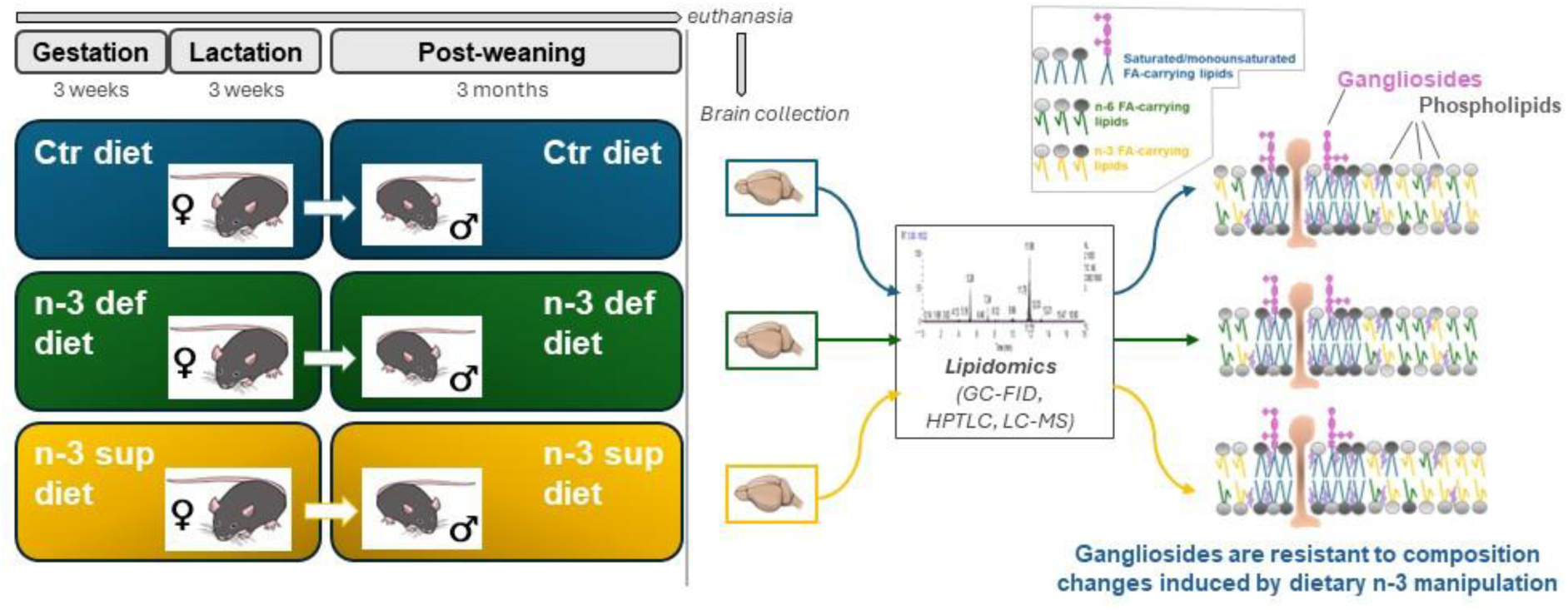

## 1 Introduction

Lipids account for no less than 50% of the dry weight of the brain, with long chain polyunsaturated fatty acids (LC-PUFAs) being of particular significance. Major brain LC-PUFAs are arachidonic acid (ARA, C20:4n-6) and docosahexaenoic acid (DHA, C22:6n-3). They are metabolites of the essential fatty acids linoleic acid (LA, C18:2n-6) and α-linolenic acid (ALA, C18:3n-3), respectively, which are the precursors of the n-6 (omega-6) and n-3 (omega-3) families. While mammals possess the enzymatic machinery to synthesize longer LC-PUFAs from precursors, the capacity of this pathway is very limited (1, 2) and n-6 and n-3 family members compete for the process. Dietary supply is therefore essential, especially during the perinatal period, and has been shown to directly impact brain content (3, 4).

LC-PUFAs are well known for their role in synaptic function, neurogenesis and neuroprotection as well as inflammation (3). Thereby, they impact brain development and function. In humans, it has been shown that adequate intake of n-3 PUFAs, especially DHA and ALA, as well as a balanced n-6/ n-3 ratio, in the mother during pregnancy and lactation is crucial for the development of the infant’s brain and visual functions (5, 6). Moreover, there is a relatively consistent body of observational studies showing that fish consumption (major dietary source of eicosapentaenoic acid (EPA) and DHA) and n-3 LC-PUFAs intake, especially EPA and DHA, are associated with a lower risk of dementia, AD and cognitive decline (7). In rodents, there are numerous studies showing the effects of diets deficient or supplemented in n-3 LC-PUFAs, especially DHA, on neurological processes, brain function and behavior. Indeed, it has been reported that chronic exposure to low levels of dietary n-3 LC-PUFAs, in particular during development, is associated with disrupted neurogenesis and neuronal differentiation, depressive- and anxiety-like behaviors in various animal models, and increased response to acute stress. On the contrary, enrichment of brain DHA levels through dietary intervention has been associated with anti-apoptotic and anti-inflammatory signaling, and increased performance in emotional and cognitive tests (3). Esterification of LC-PUFAs within the glycerophospholipids (PLs) that make up cellular membranes is one of the mechanisms thought to mediate their action in the brain since it contributes to regulate membrane biophysical properties and dynamics. Indeed, with their unsaturated acyl chains, LC-PUFAs decrease membrane bending rigidity and thus assist membrane deformation required for numerous signaling processes. Moreover, lipid-lipid and lipid-protein interactions within the plasma membrane govern localization and activity of membrane proteins, consecutively regulating their function (8). Especially, LC-PUFAs can control the dynamics of lipid rafts, which are membrane microdomains enriched in cholesterol and sphingolipids as well as functional proteins. LC-PUFAs have been shown to perturb these signaling platforms, and thus disrupt the partitioning and function of specific membrane proteins (9–11).

Beside PLs, gangliosides (GGs) are another family of complex lipids particularly abundant in plasma membrane of neurons (12). Numerous studies using neural cell cultures as well as transgenic mice deficient in specific GG biosynthetic enzymes have revealed their importance for the development, proper functioning and regeneration of the nervous system. Moreover, significant changes in brain GG profile have been reported during development and in some pathological conditions such as epilepsy, Parkinson or Alzheimer’s diseases. (12–14). GGs make a wide family of structurally diverse glycosphingolipids. They are made of a ceramide core branched with a sialylated oligosaccharide chain, both moieties exhibiting variations (15). The nature of the sugar chain, protruding toward the extracellular medium, defines the GG class and its interaction with the extracellular environment (16). The ceramide, embedded into the outer leaflet of the lipid bilayer, is composed of a long chain amino-alcohol (long chain base, LCB), mainly sphingosine (d18:1), connected to a fatty acid (FA), most of the time saturated. As PLs, GGs participate in defining the lipid composition and physicochemical properties of the plasma membrane (8). Their ability to associate laterally with each other and with sphingomyelin (SM) and cholesterol is a major driver for the formation and stabilization of membrane microdomains in which they are concentrated (17). GG biological functions rely on their presence in these lipid rafts where they interact with specific proteins leading to regulation of transmembrane signaling (18). These processes are thought to be affected by the composition of the ceramide moiety of GGs, which is tissue-specific (19) and changes with age (20) and in some pathological conditions (21–23). However, the impact of the structural diversity of the ceramide, especially changes in fatty acyl chain, on GG function is still a pending question. Little is known also about the determinants of the ceramide composition of GGs (24).

As dietary intervention using n-3 LC-PUFAs is considered a promising therapeutic approach to prevent or treat brain disorders, it is important to know the associated biochemical changes and to understand the mechanisms underlying the biological effects. While GGs are FA-carrying complex membrane lipids playing important roles in the nervous system, their contribution in mediating the effects of n-3 LC-PUFAs supply on neurological processes has not been investigated so far. The goal of the present study was to get insight into this question by determining the specific impact of dietary n-3 LC-PUFA intake on the brain GG profile and ceramide composition compared to phospholipids.

## 2 Material and methods

### 2.1 Chemicals

Chloroform (CHCl_3_) and methanol (CH_3_OH) were obtained from SDS, France. Boron trifluoride, ammonium acetate, acetonitrile (CH_3_CN), methanol and water (H_2_O) of Optima LC-MS grade were all from Fisher Scientific, France. Commercially available GG standards (GM3, GM2, GM1, GD3, GD1a, GD1b, GT1b and GQ1b) from natural sources (bovine or human) were obtained from Matreya LLC, USA. All phospholipid standards were obtained from Avanti Polar Lipids Inc., USA. Fish oil (Omegavie® 1812TG) was from Polaris (Quimper, France) and high oleic sunflower oil was from Lesieur (Coudekerque, France).

### 2.2 Animals and experimental diets

Rodent experimental conditions including dietary interventions have been described in previous publications (25, 26). Briefly, female C57BL6/J mice (Janvier Labs) were fed *ad libitum* across gestation and lactation with diets with low (control diet) or high (n-3 deficient diet) LA/ALA ratio or with a n-3 supplemented diet enriched in n-3 LC-PUFAs (stearidonic acid (SDA), EPA, DHA) (n-3 supplemented diet). The three homemade experimental diets were isocaloric (3.9 kcal/g) and isolipidic (5% fat). They contained 18.3% energy from proteins, 70.2% energy from carbohydrates and 11.5% energy from lipids They were prepared by the UPAE (Experimental Food Preparation unit, INRAE, Jouy-en-Josas, France) from a mix of synthetic lipid-free flour, whose ingredient and energy composition is given in Table 1, and oil mixtures. The oil mixtures were prepared with variable amounts of commercial sunflower oil (LA supply), canola oil (ALA supply), and fish oil (SDA, EPA and DHA supply). High oleic sunflower oil and palm oil were added to balance the levels of oleic acid and palmitic acid, respectively. The three diets differed only in their FA composition according to the oil mixture used. The complete FA composition of dietary lipids for the three different diets is given in Table 2. Offspring were maintained under the same diets after weaning and tissue collection was performed on the male offspring at adulthood (3–4-month-old).

**Table 1:** Composition of the experimental diets.

| **Ingredient** | **Amount (g/kg diet)** | **Energy (% kcal)** |
|---|---|---|
| Casein | 180 | 18.3 |
| Cornstarch | 460 | 46.8 |
| Sucrose | 230 | 23.4 |
| Cellulose | 20 |  |
| Fat<sup>1</sup> | 50 | 11.5 |
| Mineral mix<sup>2</sup> | 50 |  |
| Vitamin mix<sup>3</sup> | 10 |  |

<sup>1</sup>represented by oil mixtures prepared with variable amounts of sunflower oil, canola oil, high oleic sunflower oil, palm oil and fish oil depending on the experimental diet (composition in Table 2)
<sup>2</sup>composition (g/kg): sucrose, 110.7; CaCO<sub>3</sub>, 240; K<sub>2</sub>HPO<sub>4</sub>, 215; CaHPO<sub>4</sub>, 215; MgSO<sub>4</sub>,7H<sub>2</sub>O, 100; NaCl, 60; MgO, 40; FeSO<sub>4</sub>,7H<sub>2</sub>O, 8; ZnSO<sub>4</sub>,7H<sub>2</sub>O, 7; MnSO<sub>4</sub>,H<sub>2</sub>O, 2; CuSO<sub>4</sub>,5H<sub>2</sub>O, 1; Na<sub>2</sub>SiO<sub>7</sub>,3H<sub>2</sub>O, 0.5; AlK(SO<sub>4</sub>)<sub>2</sub>,12H<sub>2</sub>O, 0.2; K<sub>2</sub>CrO<sub>4</sub>, 0.15; NaF, 0.1; NiSO<sub>4</sub>,6H<sub>2</sub>O, 0.1; H<sub>2</sub>BO<sub>3</sub>, O.1; CoSO<sub>4</sub>,7H<sub>2</sub>O, 0.05; KIO<sub>3</sub>, 0.04; (NH<sub>4</sub>)6Mo<sub>7</sub>O<sub>24</sub>,4H<sub>2</sub>O, 0.02; LiCl, 0.015; Na<sub>2</sub>SeO<sub>3</sub>, 0.015; NH<sub>4</sub>VO<sub>3</sub>, 0.01
<sup>3</sup>composition (g/kg): sucrose, 549.45; retinyl acetate, 1; cholecalciferol, 0.25; DL-a-tocopheryl acetate, 20; phylloquinone, 0.1; thiamin HCl, 1; riboflavin, 1; nicotinic acid, 5; calcium pantothenate, 2.5; pyridoxine HCl, 1; biotin, 1; folic acid, 0.2; cyanobalamin, 2.5; choline HCl, 200; DL-methionin, 200; p-aminobenzoic acid, 5; inositol, 10

**Table 2.** Fatty acid composition of dietary lipids (% of total FAs)

| **Diets** | **Control (balanced)** | **n-3 PUFA-deficient** | **n-3 PUFA-supplemented** |
|---|---|---|---|
| C16:0 | 22.6 | 6.2 | 20.7 |
| C18:0 | 3.3 | 4.4 | 3.6 |
| Other saturated FAs | 1.8 | 1.6 | 5.9 |
| Total saturated FAs | 27.7 | 12.2 | 30.2 |
| C16:1n-7 | 0.2 | 0.1 | 5.3 |
| C18:1n-9 | 57.9 | 26 | 22.6 |
| C18:1n-7 | 1.5 | 0.9 | 4.9 |
| Other monounsaturated FAs | 0.4 | 0.2 | 1.3 |
| Total monounsaturated FAs | 60 | 27.2 | 34.1 |
| C18:2n-6 (LA) | 10.6 | 60.5 | 15.1 |
| C20:4n-6 (ARA) | ND | ND | 0.6 |
| Total n-6 LC-PUFAs | 10.7 | 60.5 | 15.7 |
| C18:3n-3 (ALA) | 1.6 | 0.1 | 1.1 |
| C18:4n-3 (SDA) | ND | ND | 2.1 |
| C20:5n-3 (EPA) | ND | ND | 10.3 |
| C22:5n-3 (n-3DPA) | ND | ND | 0.7 |
| C22:6n-3 (DHA) | ND | ND | 5.8 |
| Total n-3 LC-PUFAs | 1.6 | 0.1 | 20 |
| Total PUFAs | 12.3 | 60.6 | 35.7 |
| n-6/n-3 | 6.7 | 605 | 0.8 |

Lipids in the diets were provided by specific oil mixtures composed of 3.2% sunflower oil, 32.5% high oleic sunflower oil, 43.0% palm oil and 21.3% canola oil for the control diet; 100% sunflower oil for the n-3 PUFA-deficient; 14.2% sunflower oil, 1.3% high oleic sunflower oil, 3.8% palm oil, 15.4% canola oil and 65.3% fish oil for the n-3 PUFA-supplemented diet.
FAs, fatty acids; LA, linoleic acid; ARA, Arachidonic acid; LC-PUFAs, long chain polyunsaturated fatty acids; ALA, α-linolenic acid; SDA, stearidonic acid; EPA, eicosapentaenoic acid; DPA, docosapentaenoic acid; DHA, docosahexaenoic acid.
Values are expressed as percentages of total FAs. ND, not detected.

### 2.3 Tissue sample preparation and lipid extraction

First, mouse half-brains were homogenized in 0.5 ml H_2_O using tungsten microbeads and a tissue lyser (Qiagen, The Netherlands) during 1 min 30 s at 30 Hz speed. Total lipids were extracted with 5 mL of CHCl_3_/CH_3_OH (1:1, v/v), overnight at 4°C. The residual pellet obtained after centrifugation (1,500 *g*, 5 min) was reextracted with 3 mL CHCl_3_/CH_3_OH (1:1, v/v), 3 mL CHCl_3_/CH_3_OH (2:1, v/v) and 3 mL CHCl_3_/CH_3_OH/H_2_O (48:35:10,v/v/v). The four lipid extracts were combined and dried under a stream of nitrogen. Total lipids were weighed then re-dissolved in CHCl_3_/CH_3_OH (1:1, v/v) for the different analyses described below.

### 2.4 Fatty acid analysis by gas chromatography coupled to a flame ionization detector (GC-FID)

Total lipid extracts (equivalent to 2 mg total lipids) were analyzed using GC-FID after transmethylation as previously described (27). Lipid extracts were transmethylated using boron trifluoride in methanol for 2 hours at 95°C, producing fatty acid methyl esters (FAMEs) from acyl-moieties of conventional PLs and plasmalogens, and dimethylacetals (DMAs) from alkenyl-moieties of plasmalogens (28). Both were subsequently extracted with hexane and analyzed by gas chromatography on a GC Trace 1310 gas chromatograph (Thermo Scientific, Les Ulis, France) using a CPSIL-88 column (100mx0.25mm i.d., film thickness 0.20µm; Varian, Les Ulis, France) equipped with a flame ionization detector. Hydrogen was used as the carrier gas (inlet pressure 210kPa). The oven temperature was held at 60°C for 5 min, increased to 165°C at 15°C/min and held for 1 min, and then to 225°C at 2°C/min and finally held at 225°C for 17 min. The injector and the detector were maintained at 250°C. FAMEs and DMAs were identified by comparison with commercial and synthetic standards. The data were processed using the EZChrom Elite software (Agilent Technologies, Massy, France) and reported as a percentage of the total signal intensity corresponding to the sum of all detected FAMEs and DMAs.

### 2.5 Phospholipid analyses

PL analyses were performed as described in (29, 30). Methods are briefly presented below.

#### 2.5.1 Separation and relative quantification of phospholipid classes using liquid chromatography (LC) coupled to charged aerosol detection (Corona-CAD)

After measurement of the phosphorus content of the total lipid extracts (31), samples were diluted to 500 μg/mL PLs in CHCl_3_/CH_3_OH 1:1 (v/v) for analysis. PL classes were separated by high-performance liquid chromatography (HPLC) under hydrophilic interaction liquid chromatography (HILIC) conditions according to their increasing polarities. PL classes were detected and quantified using a Corona™ universal detector performing charged aerosol detection (Ultra RS Charged Aerosol Detector™ CCAD, Thermo Scientific, USA). The data were reported as a percentage of the total signal intensity corresponding to the sum of all detected PL classes.

#### 2.5.2 Separation, characterization and relative targeted quantification of phospholipid species by liquid chromatography coupled to mass spectrometry (LC-MS)

PL classes were separated by HPLC under HILIC conditions. For characterization of PL molecular species, the chromatographic system was coupled to an Orbitrap Fusion™ Tribrid mass spectrometer (Thermo Scientific, USA) equipped with an EASY-Max NGTM ion source (heated electrospray ionization H-ESI). PL molecular species were characterized based on high-accuracy MS data and MS/MS experiments, performed alternatively in positive and negative mode, using the LipidSearch™ software and LIPID MAPS™ database. For targeted quantification, the chromatographic system was coupled to a triple quadrupole mass spectrometer (QqQ) (Thermo Finnigan TSQ Quantum) equipped with a standard electrospray ionization source. Phosphatidylethanolamine (PE) species were quantified by neutral loss scanning of 141 Da, which corresponds to the ethanolamine phosphate head, in the positive ion mode by precursor ion scanning. Phosphatidylcholine (PC) and sphingomyelin (SM) species were quantified in the positive ion mode by precursor ion scanning of *m/z* 184 amu, which corresponds to the inositol phosphate head group (minus H_2_O). The data were processed using the Xcalibur software. Corrections were applied to the data for isotopic overlap. The level of each PL molecular species was expressed relative to the sum of all quantified species in its specific PL class.

### 2.6 Ganglioside analyses

#### 2.6.1 Ganglioside extraction and purification

Total lipid extracts (equivalent to 4.5 mg total lipids) were dried under a stream of nitrogen, and redissolved in CHCl_3_/CH_3_OH (1:1, v/v). GGs were then separated from other lipids by phase partition by adding H_2_O. After centrifugation, the upper aqueous phase was collected while the lower organic phase was reextracted twice with CH_3_OH/water (1:1, v/v). The three upper phases containing GGs were combined, dried under a stream of nitrogen, and redissolved in CH_3_OH/PBS 10 mM (1:1, v/v) in order to be purified by solid phase extraction using C18 silica gel columns (Sep-Pak Vac 6cc, 500 mg; Waters). GGs were first washed with H_2_O and purified GGs were eluted with CH_3_OH and CHCl_3_/CH_3_OH (2:1, v/v). This procedure is described in further details in (32).

#### 2.6.2 GG-bound sialic acid (GG-SA) measurement

GG-SA was measured on purified GG extracts (equivalent to 0.7 mg total lipids) using the procedure of Svennerholm with slight modifications (33). Commercial SA (Sigma-Aldrich) was used as a standard for the assay and resorcinol reagent was used to detect SA after heating at 110°C for 20 min, as previously described (32). Optical density was finally read at 580 nm in a spectrophotometer.

#### 2.6.3 Quantification of ganglioside classes using High Performance Thin Layer Chromatography (HPTLC)

Aliquots of the purified GG extracts (equivalent to 10 nmol GG-bound sialic acid) were applied to a HPTLC plate (Merck, Germany). The plate was developed in CHCl_3_/CH_3_OH/0.2% CaCl_2_ (55:45:10, v/v/v) and revealed with resorcinol reagent, which detects GG-SA, as described in (32). GGs, appearing as purple-blue bands, were identified by co-migration with standards. A standard curve was also performed for each GG class to quantify GGs by densitometric analysis using a CCD camera (Chemidoc™ MP System from Biorad, USA). The distribution of GG classes was expressed as a percentage of the total nmol of GG detected.

#### 2.6.4 Fatty acid analysis by GC-FID

Total lipid extracts (equivalent to 2 mg total lipids) were subjected to alkaline hydrolysis using KOH 1M in MeOH for 2h at 37°C to hydrolyze PLs. GGs were then separated by phase partition as described in 2.6.1, except that water was acidified (pH=2) to avoid recuperation of FAs hydrolyzed from PLs along with GGs. FAMEs were then produced and analyzed by GC-FID as described in section 2.4 except trans-methylation was performed overnight.

#### 2.6.5 Relative targeted quantification of ganglioside species by LC-MS

Purified GG extracts were analyzed using MS after prior separation using a method of liquid chromatography under specific HILIC conditions, as previously described (32). The QqQ mass spectrometer (Thermo Finnigan TSQ Quantum) was operated in negative ion mode. GGs produced an abundant product ion at *m/z* 290 amu corresponding to a characteristic *N*-Acetylneuraminic acid (sialic acid) fragment. GG classes and molecular species were identified by comparison with commercial standards when available and/or based on the theoretical mass and previous identification of brain GGs performed in humans using high-resolution LTQ_Orbitrap XL mass spectrometer and MS/MS (19). For quantification, data were acquired in Selected Reaction Monitoring (SRM). Molecular species carrying PUFAs were specifically looked for (18:3, 20:4, 20:5, 22:4, 22:5 and 22:6). The proportion of each ceramide species in a specific GG class was calculated as the ratio of its peak area to the sum of all detected peak areas in this class, every GG class being considered separately.

### 2.7 Statistical analyses

Results were submitted to non-parametric statistical analyses using GraphPad Prism 10 software (GraphPad Software, LLC). Mann-Whitney tests were performed to independently compare the deficient and supplemented groups to the control group. *p<0.05, **p<0.005

## 3 Results

### 3.1 The fatty acid composition of the brain is modified by the diet

The analysis of total lipids performed using gas chromatography after trans-methylation gave an overview of the relative abundance of FAs carried mainly by PLs since they represent the large majority of brain lipids. The complete FA composition of mouse brain under the different diets is available in Supplementary table 1 and a graphic representation of the main FA species is given in Figure 1A. As expected, brain lipid profile is characterized by a significant proportion of LC-PUFAs (nearly 27% of total fatty acids) and high levels of DHA, which accounts for 13% of total FAs. The levels of LC-PUFAs from the n-6 and n-3 families were substantially affected by the experimental diets. The n-3 deficient diet led to increased levels of FAs of the n-6 family and decreased levels of FAs of the n-3 family compared to the control diet. Especially, the levels of C20:4n-6 (ARA), C22:4n-6 and C22:5n-6 (n-6 DPA) were increased while those of C22:5n-3 (n-3 DPA) and C22:6n-3 (DHA) were decreased. On the contrary, the n-3 supplemented diet led to decreased levels of FAs of the n-6 family and increased levels of FAs of the n-3 family compared to the control diet. Especially, the levels of ARA, C22:4n-6 and n-6 DPA were decreased while those of C20:5n-3 (EPA), undetected under control conditions, and those of n-3 DPA and DHA were increased. These modifications strongly impacted the n-6/n-3 ratio which was multiplied 2.7-fold by the n-3 deficient diet and divided 2-fold by the n-3 supplemented diet compared to the control diet (Figure 1B). The level of other FAs was also modified by the diet such as oleic acid (C18:1n-9) which was decreased by the n-3 deficient diet and increased by the n-3 supplemented diet.

**Figure 1.**
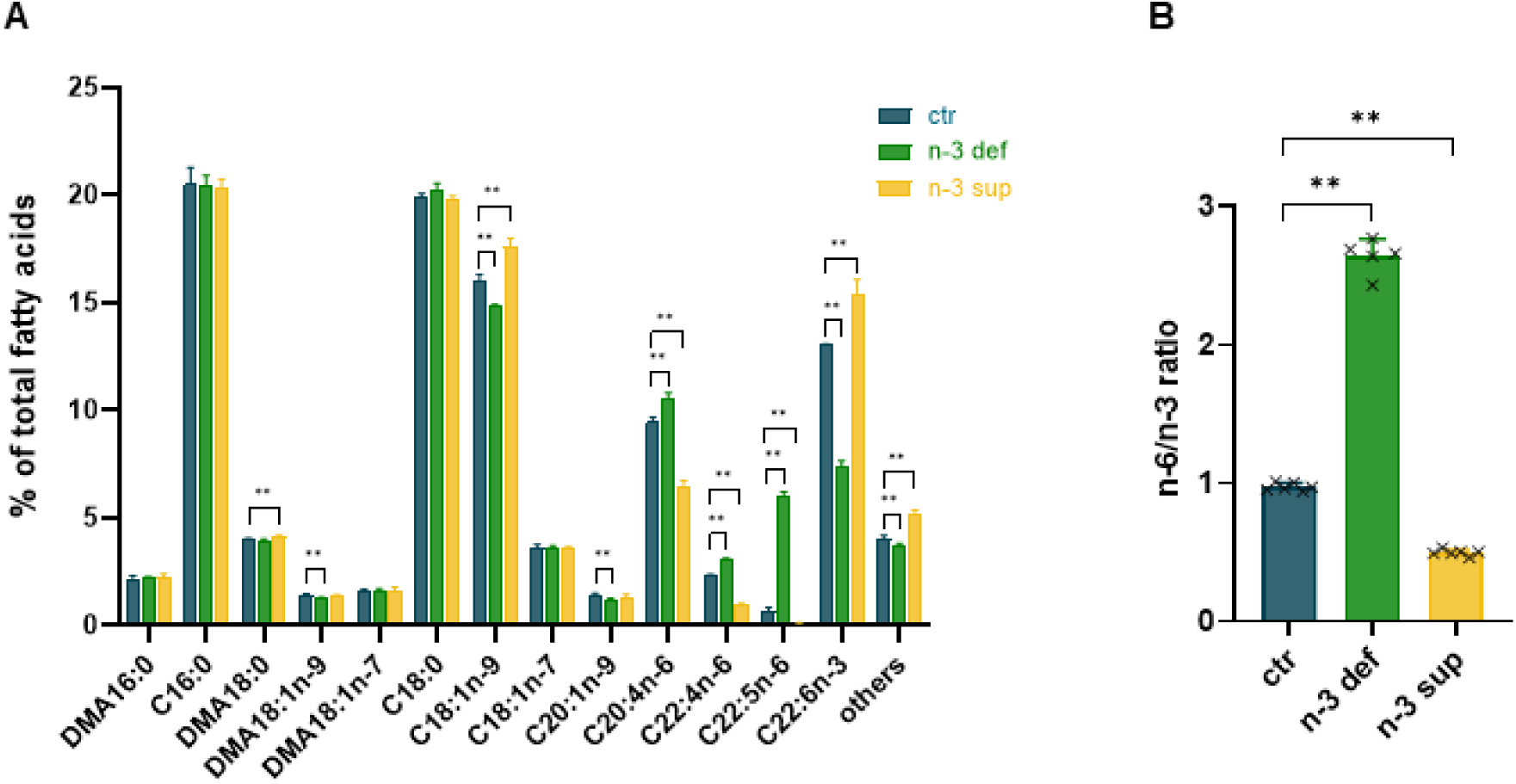
Fatty acid composition of the mouse brain. Brain total lipids of mice subjected to the different diets were analyzed using gas chromatography coupled to mass spectrometry (GC-FID) after trans-methylation. (A) The fatty acid (FA) profile is represented by the relative level of each detected FA expressed as a percentage of total FAs (FA methyl esters (FAMEs) and dimethylacetals (DMAs)). They are abbreviated as number of carbons:number of double bonds. FAs/DMAs which accounted for less than 1% in all three diet conditions were grouped under “others”. (B) The n-6/n-3 ratio was calculated by dividing the sum of the percentages of all detected n-6 FAs by the sum of the percentages of all detected n-3 FAs. Each bar represents the mean ± SD of 5-6 mice. Individual data are shown. ** p< 0.005.

### 3.2 The brain phospholipid profile is unaffected by the diet

The distribution of the different PL classes determined using liquid chromatography coupled to a universal Corona detector is presented in Figure 2. PI, PE, PC, lyso-PC (LPC) and SM were quantified. PE and PC were by far the most abundant, accounting for about 93% of total PLs. We observed that the distribution of the PL classes did not change in response to the diet, except for SM whose relative levels were slightly decreased in response to the n-3 supplemented diet.

**Figure 2.**
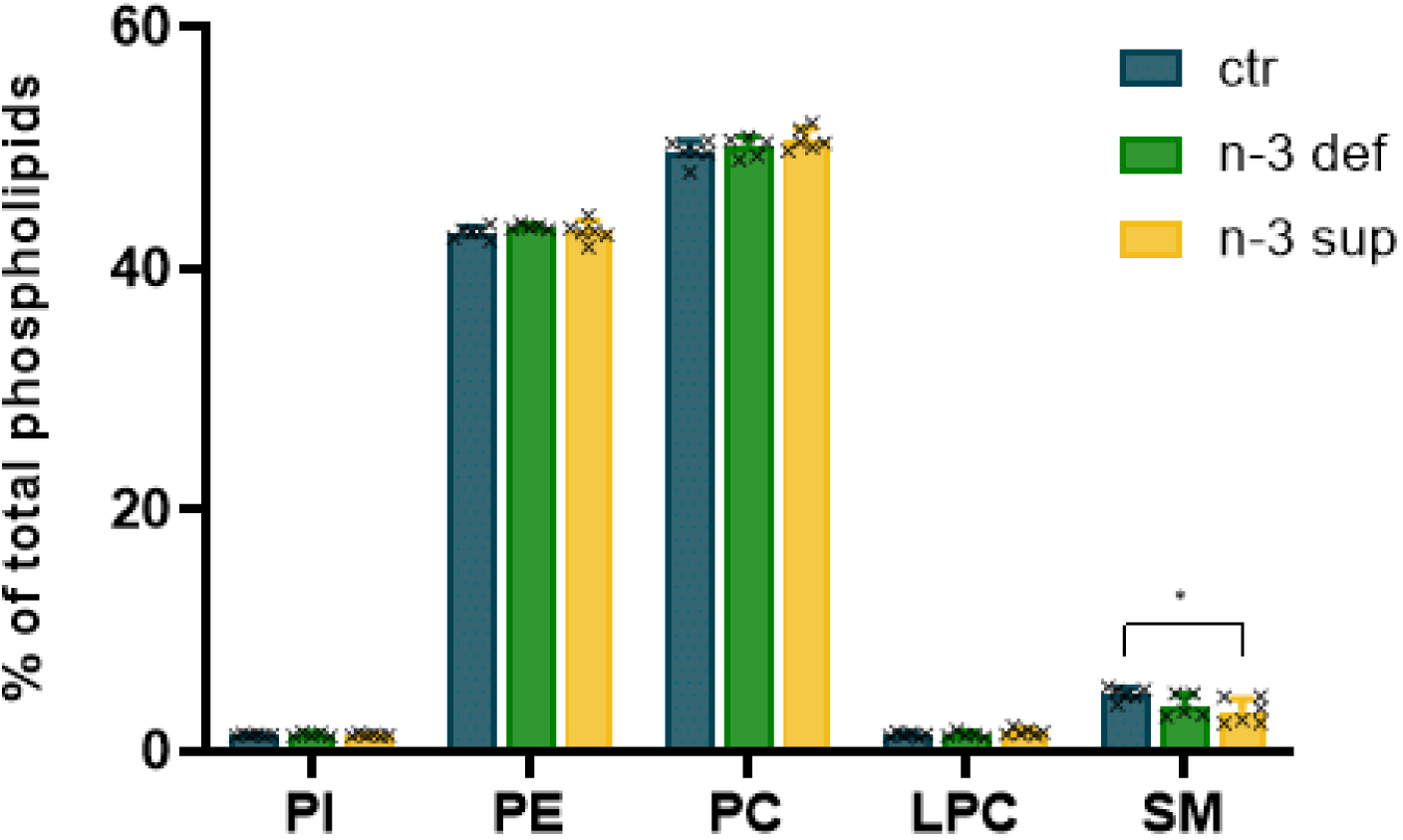
Phospholipid profile of the mouse brain. Brain total lipids of mice subjected to the different diets were analyzed using high-performance liquid chromatography (HPLC) coupled to a Corona detector. The phospholipid (PL) profile is represented by the relative level of each PL class expressed as a percentage of total PLs. Each bar represents the mean ± SD of 5-6 mice. Individual data are shown. *p<0.05. PI, Phosphatidylinositol; PE, phosphatidylethanolamine; PC, phosphatidylcholine; LPC, lyso-PC; SM, sphingomyelin.

### 3.3 The distribution of individual species of brain glycerophospholipids changes in response to the diet

The distribution of the individual species of the two major PL classes, PEs and PCs, was determined using LC-MS. Identification of the FAs carried by the PLs was performed using high resolution MS and quantification was performed using a triple quadrupole. The complete distribution of phospholipid species under the different diets is available in Supplementary tables 2 and 3 and a graphic representation of the main species is given in Figure 3A for PEs and in Figure 3B for PCs. Results show that this distribution is specific to the class considered. Indeed, the most abundant species within PEs are PE 18:0/22:6 and PE 18:0/20:4 while they are PC 16:0/18:1, PC 16:0/16:0, and PC 18:0/18:1 in PCs. We observed that the two experimental diets were associated with changes in the distribution of individual species. The relative abundance of PE and PC molecular species carrying DHA were decreased in response to the n-3 deficient diet and increased in response to the n-3 supplemented diet. On the contrary, the relative abundance of PE and PC molecular species carrying ARA, 22:5 or 22:4 FAs were increased in response to the n-3 deficient diet and decreased in response to the n-3 supplemented diet. PE 18:1/18:1 levels also increased in response to the n-3 deficient diet and decreased in response to the n-3 supplemented diet. Overall, those results are in accordance with the FA profile data (Figure 1). Of note, LC-MS does not allow to distinguish between n-6 and n-3 families. Whereas 22:5 FA does exist in both families, the FA composition obtained using GC-FID (Supplementary table 1) indicates that the levels of 22:5n-6 are much higher than those of 22:5n-3, thus suggesting that the 22:5 FA carried by PLs detected using LC-MS is most probably n-6 DPA.

**Figure 3.**
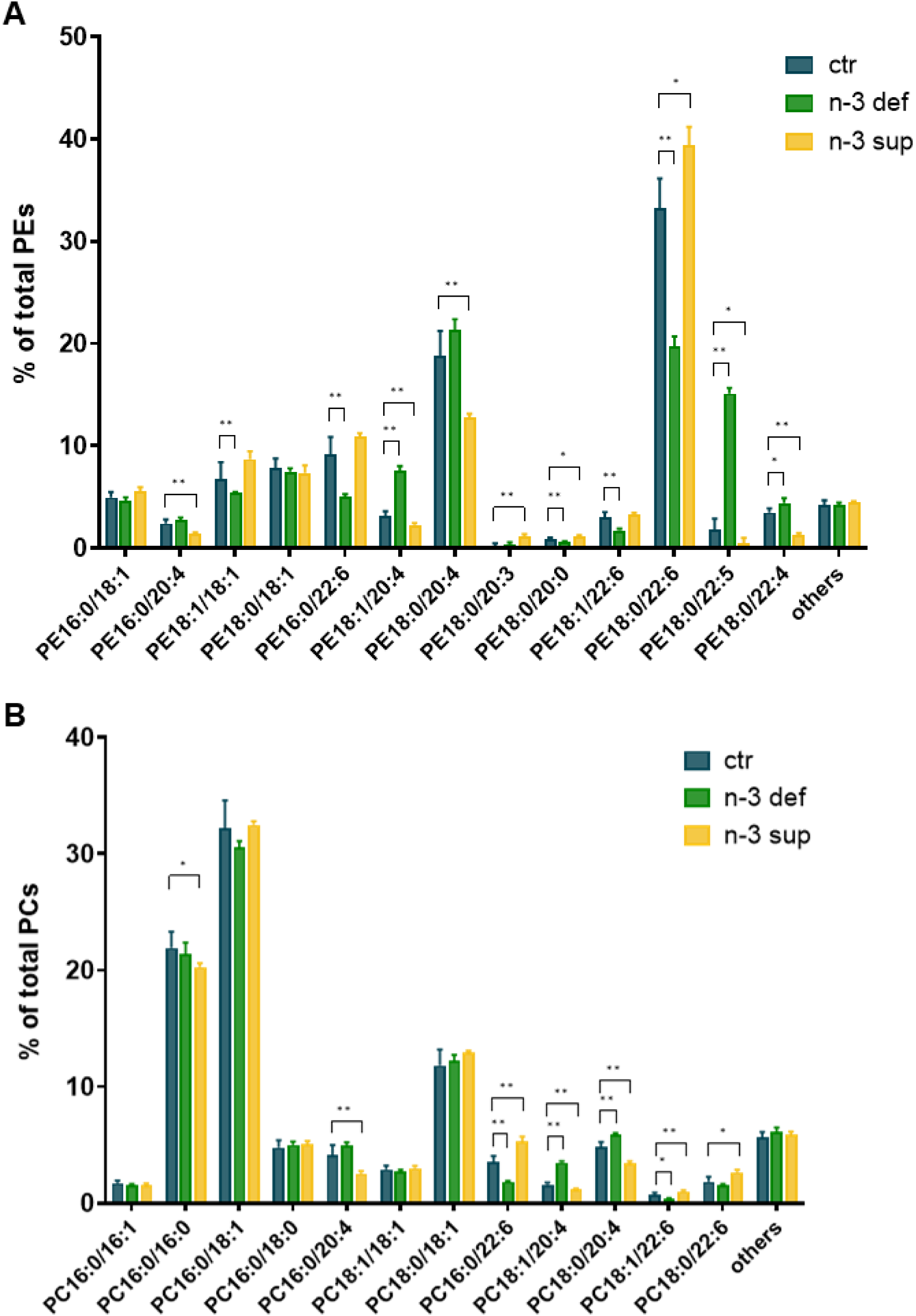
Distribution of the individual species of mouse brain glycerophospholipids. Brain phosphatidylethanolamines (PEs) (A) and phosphatidylcholines (PCs) (B) of mice subjected to the different diets were analyzed using liquid chromatography coupled to mass spectrometry (LC-MS). Quantification of the molecular species was performed using a triple quadrupole and identification using a Orbitrap Fusion. Position on the glycerol backbone is shown as sn-1/sn-2 of the fatty acid radicals. A minor isobaric species was detected for PE 18:1/20:4 (PE 16:0/22:5), PE 18:0/20:4 (PE 16:0/22:4, traces), PC 16:0/16:1 (PC 14:0/18:1), PC 18:0/18:1 (PC 16:0/20:1, traces). The specific profile is represented by the relative level of each detected molecular species expressed as a percentage of the sum of all molecular species signal intensities in a specific PL class. Molecular species which accounted for less than 1% in all three diet conditions were grouped under “others”. Each bar represents the mean ± SD of 5-6 mice. *p<0.05, **p< 0.005.

### 3.4 The distribution of individual species of brain sphingomyelins is not modified by the diet

The distribution of the individual SM species was determined using LC-MS. Identification of the FA and LCB carried by the SMs was performed using high resolution MS and quantification was performed using a triple quadrupole. The complete distribution of ceramide species under the different diets is available in Supplementary table 4 and a graphic representation is given in Figure 4 for the main species. The LCB were exclusively d18:1, apart from 2 molecular species carrying d18:2 and detected as traces amount (Sup table 4). The major molecular species was d18:1/18:0 (about 70% of all quantified species) followed by d18:1/24:1 (about 12% of all quantified species). It appears that SMs, contrary to PLs, contain only saturated and mono-unsaturated FAs, and no PUFAs were detected. Apart from a slight increase of the levels of d18:1/18:1 species under the n-3 deficient diet, there was no modification of the SM species relative abundance in response to the diets.

**Figure 4.**
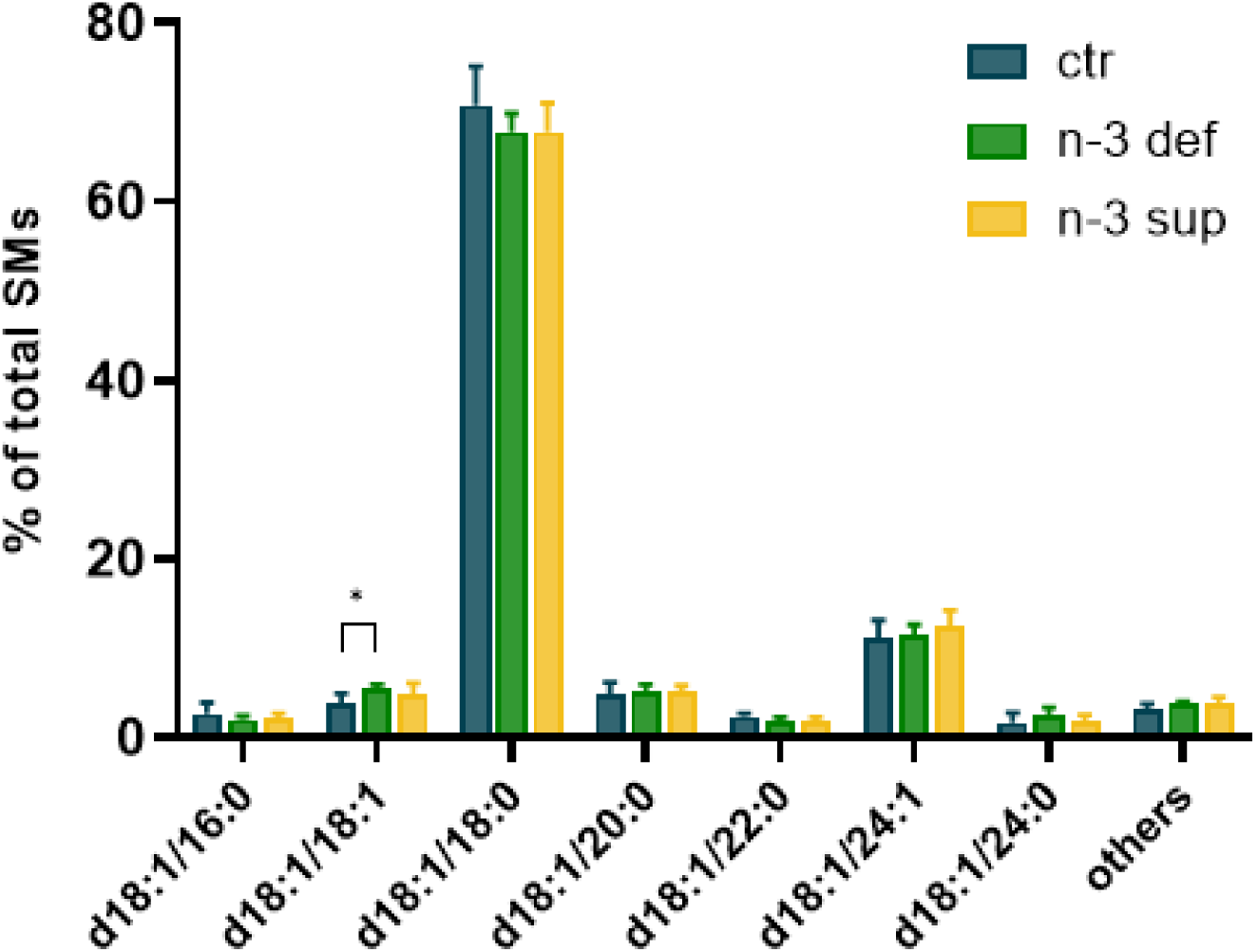
Distribution of the individual species of mouse brain sphingomyelins. Brain sphingomyelins (SMs) of mice subjected to the different diets were analyzed using liquid chromatography coupled to mass spectrometry (LC-MS). Quantification of the molecular species was performed using a triple quadrupole and identification using a Orbitrap Fusion. Position on the ceramide backbone is shown as long chain base/fatty acid radical. The specific profile is represented by the relative level of each detected molecular species expressed as a percentage of the sum of all molecular species signal intensities. Molecular species which accounted for less than 1% in all three diet conditions were grouped under “others”. Each bar represents the mean ± SD of 5-6 mice. *p<0.05.

### 3.5 The ganglioside profile of the brain is not affected by the diet

The GG-bound sialic acid (GG-SA) content, measured using colorimetry as described under Material and Methods, was used as an estimate of the total GG content. As shown in Figure 5A, mouse brain contained an average of 19.6 ± 2.47 nmol GG-SA / mg lipids and the different experimental diets had no significant effect on this amount. The distribution of the different GG classes was studied by HPTLC and colorimetric revelation (Figure 5B). GG classes were identified by co-migration with standards. Mouse brain contained mainly polysialo-GGs. The major ones were GD1a and GT1b, accounting together for about 60% of all GGs, followed by GM1, GD1b, and GQ1b, accounting for 12%, 9% and 14% of all GGs, respectively. GD2 and GD3 were present as minor classes. Traces of GM3 could also be detected. Semi-quantitative results presented in Figure 5C indicated no change of GG profile in response to the different diets.

**Figure 5.**
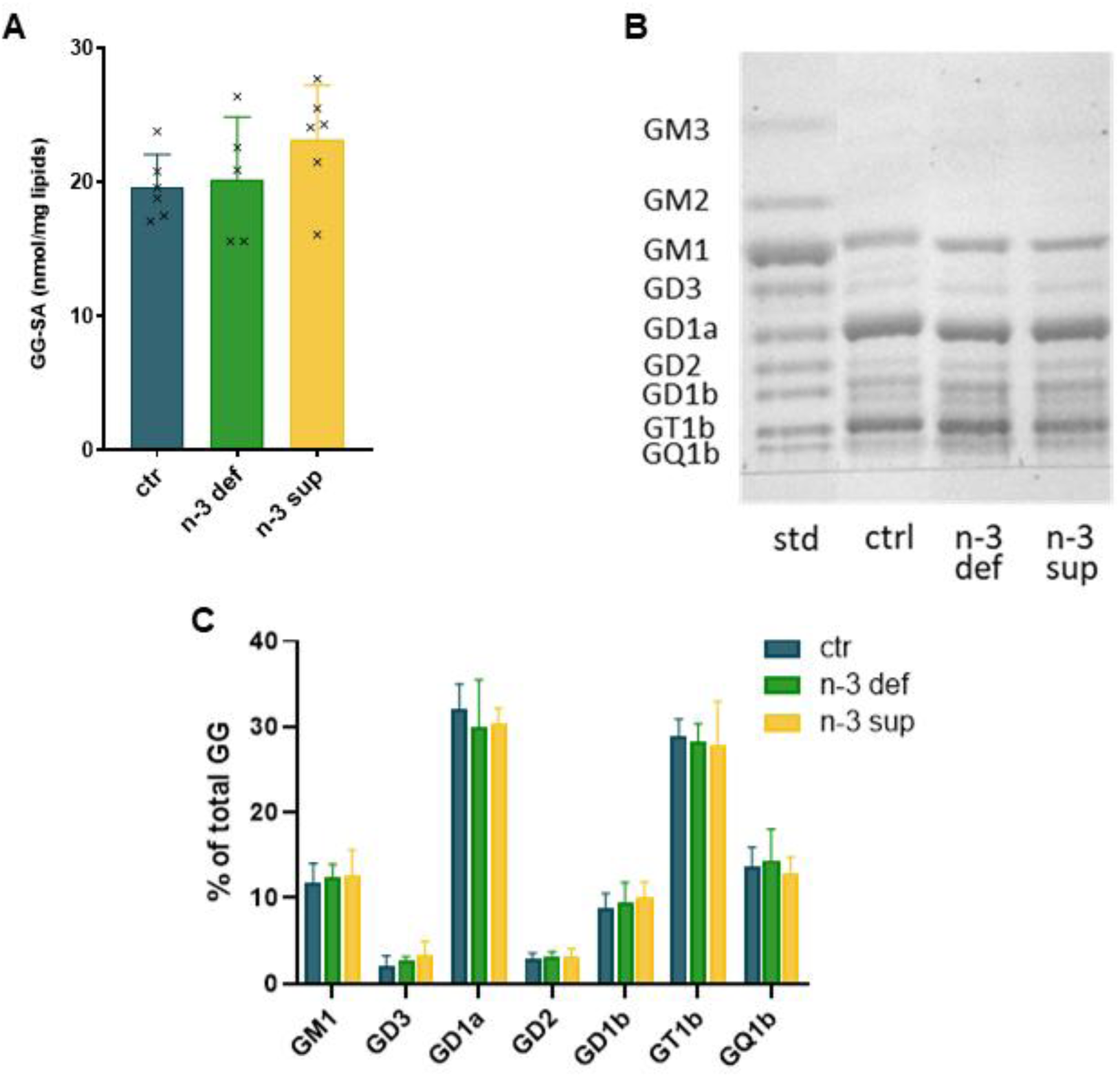
**Ganglioside profile of the mouse brain**(A) Total GG-SA (Ganglioside-bound sialic acid) was quantified on GG extracted from the brain of mice subjected to the different diets. Each bar represents the mean ± SD of 5-6 mice. Individual data are shown. GG classes were analyzed using HPTLC (high performance thin layer chromatography). (B) A representative profile is shown for each diet condition (10 nmol LBSA spotted) as well as a mixture of GG standards (1 nmol LBSA each). (C) Using standard curves and densitometric analysis, detected GG classes were quantified and their relative distribution, expressed as percentage of total nmol of GGs, was calculated. Each bar represents the mean ± SD of 5-6 mice.

### 3.6 The fatty acid composition of brain gangliosides is not modified by the diet

After purification of brain GGs, FAs were analyzed following trans-methylation using GC-FID. The complete distribution of GG FAs under the different diets is available in Supplementary table 5 and a graphic representation of the main species is given in Figure 6. C18:0 represented by far the major FA of brain GGs, accounting for almost 90% of all GG FAs. The other ones were all saturated FAs, except for C18:1n-9. Also, C16:1n-9, C18:2n-6 (LA) and C24:0 could be detected but were grouped under “others” since they represented less than 1% of all FAs. No LC-PUFA was detected and the FA profile of brain GG was similar in every diet condition.

**Figure 6.**
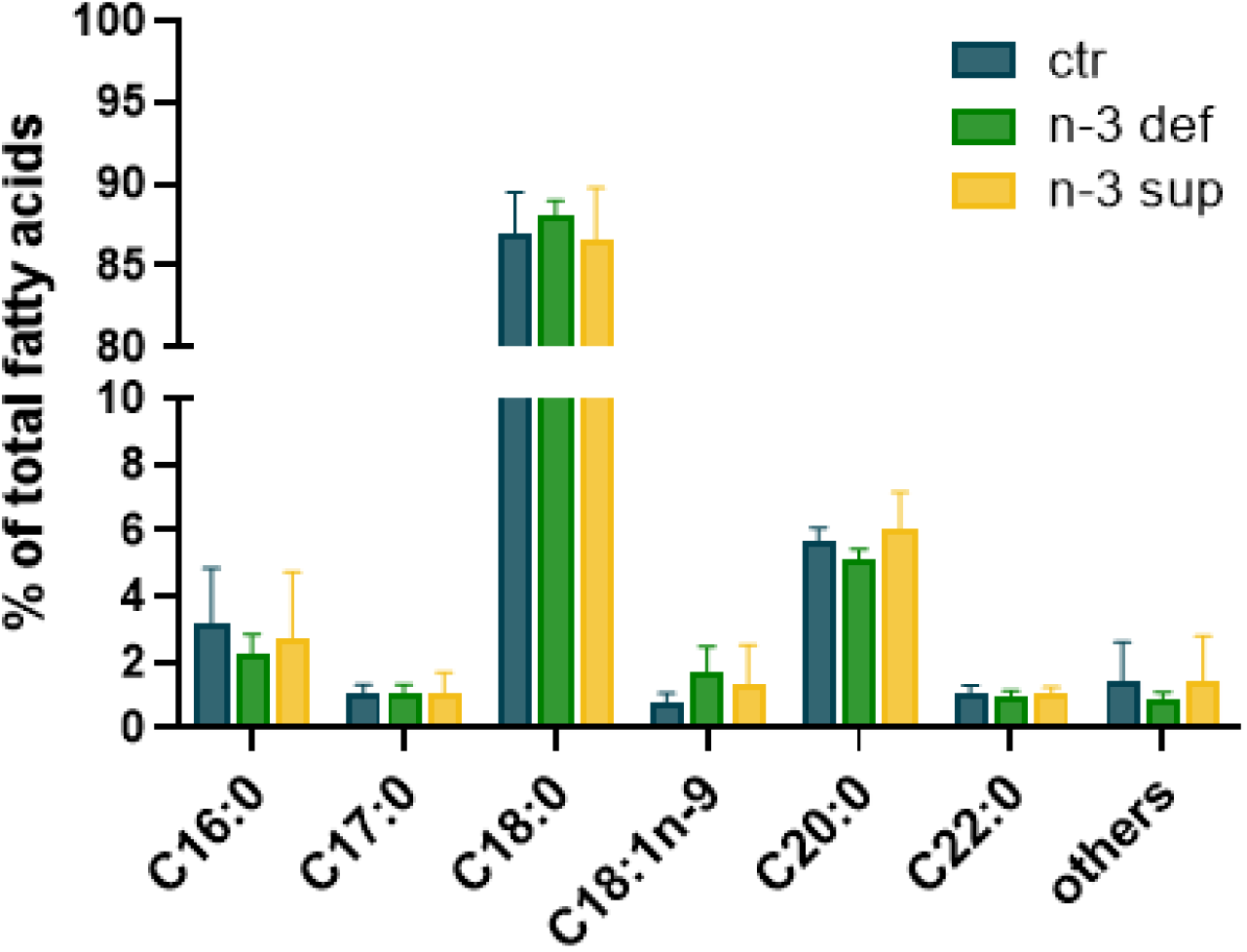
Fatty acid composition of mouse brain gangliosides. After hydrolysis of phospholipids and extraction of gangliosides (GGs) from total lipid extracts, fatty acids (FAs) of the brain GGs of mice subjected to the different diets were analyzed using GC-FID (gas chromatography coupled to a flame ionization detector) after trans-methylation. The FA profile is represented by the relative level of each detected FA expressed as a percentage of the sum of all FA signal intensities. FAs which accounted for less than 1% in all experimental conditions were grouped under “others”. Each bar represents the mean ± SD of 5-6 mice.

### 3.7 The distribution of the individual species of brain gangliosides does not change in response to the diet

GGs were further studied using MS after prior separation of the GG classes using a method of liquid chromatography. Figure 7 shows representative ion chromatograms of the mouse brain GGs of the three experimental groups. The GG classes previously identified using HPTLC and colorimetric revelation were detected. In addition, minor GG classes could be detected thanks to the sensitivity of the LC-MS technique: GM4, a gala-series GG, as well as GM4 carrying a 2-hydroxy FA, GM3, GM2, and the acetylated forms of GT1b and GQ1b. Of note, GM1 could not be detected with the triple quadrupole mass spectrometer because it did not form multicharged ions and therefore stand over the mass range of the apparatus. It should be noted that this LC-MS method is not quantitative since the peak intensity of a GG class is not directly proportional to its absolute amount but also depends on the ability of this specific class to be fragmented and ionized in the apparatus. Nevertheless, a specific GG class can be compared between the different experimental groups. It appears that the distribution of the different GG classes is not significantly affected by the diet.

**Figure 7.**
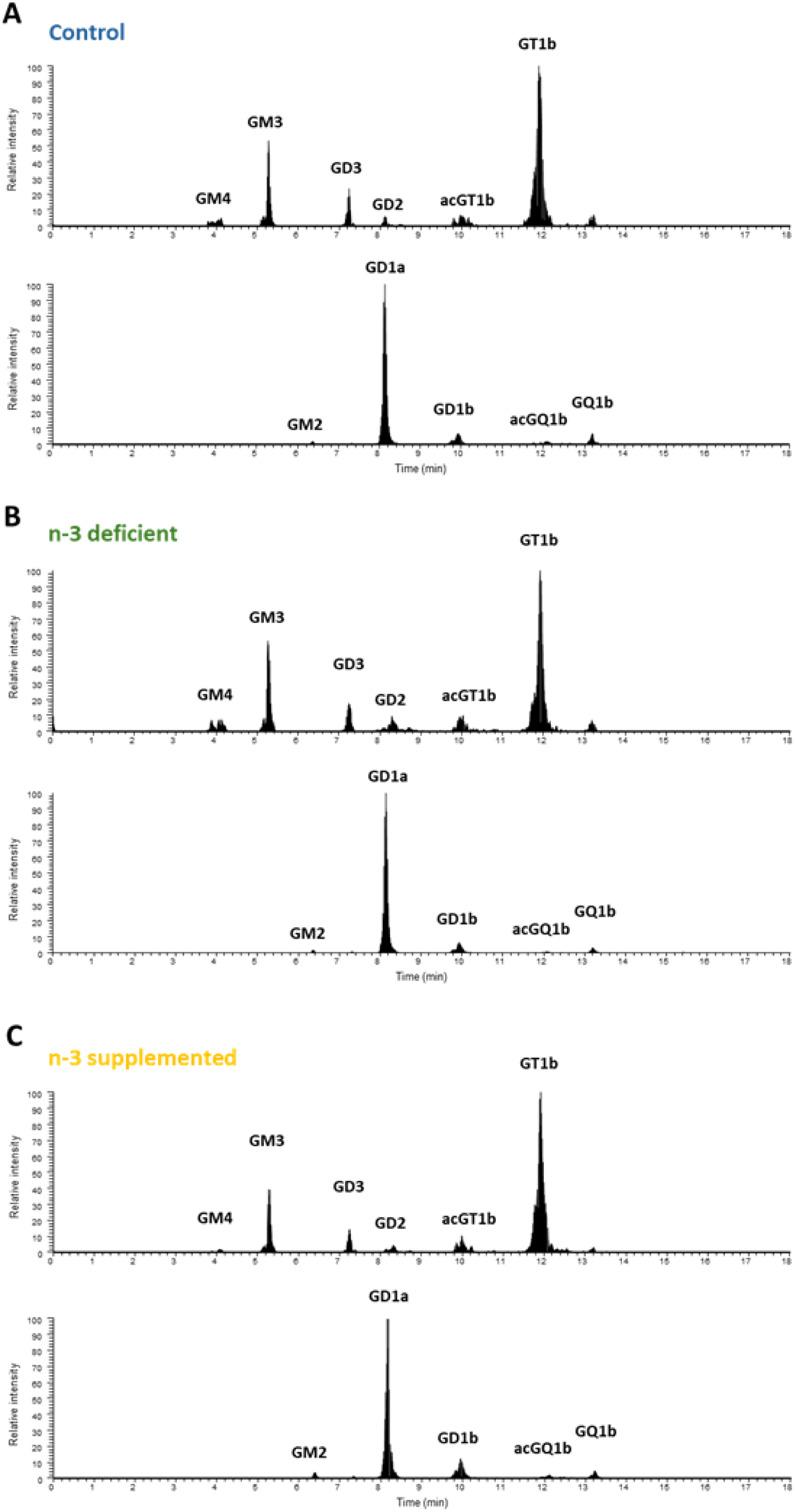
Liquid chromatography coupled to mass spectrometry (LC-MS) ganglioside profile of the mouse brain. Brain gangliosides (GGs) of mice subjected to the different diets were analyzed using LC-MS. Representative precursor ion chromatograms of the GGs of mouse brain under control (A), deficient (B) and n-3 supplemented diet (C) are shown. The QqQ mass spectrometer was operated in the precursor ion scanning of the characteristic fragment of GGs, N-Acetylneuraminic acid, at m/z 290, in the negative ionization mode. GG classes were identified by comparison with commercial standards when available, with theoretical mass and based on previous identification of brain GGs performed in humans using high-resolution LTQ_Orbitrap XL mass spectrometer (19). To optimize detection of every GGs, GG classes were distributed into two distinct analytical methods for each sample. Results are presented as relative signal intensities for each method. GM1, which does not form multicharged ions and stand over the mass range of the spectrometer could not be detected.

MS allows for the detection of every molecular species present in each GG class, based on their exact mass and charge state. They correspond to the different ceramide moieties made of a LCB and a FA both varying in length and number of unsaturations. A large diversity of ceramide species was detected and quantified among the different GG classes. The proportion of each species was determined in the different GG classes considered independently. The complete ceramide distribution of GGs under the different diets is available in Supplementary table 6 and a graphic representation of the main species is given in Figure 8. As expected, GGs are mainly made of monounsaturated ceramides. The most abundant ceramide species in most GG classes is 36:1, accounting for 57 to 87 % of all quantified species under control conditions. Ceramide 38:1 comes next accounting for 8 to 30 %. Since sphingosine (d18:1) and d20:1 have been reported as the most abundant LCBs in brain GGs (19, 34–36), these results are in accordance with our analysis of the FA profile of brain GGs (Fig. 6), which showed that 18:0 accounts for almost 90% of all FAs. GM4 and GM4OH, which do not belong to the ganglio-series GGs, make exception by exhibiting a peculiar ceramide profile dominated by 42:2, 42:1 and 40:1 ceramides. Overall, it appeared that the distribution of the GG ceramide species within the GG classes was very similar between the different diets.

**Figure 8.**
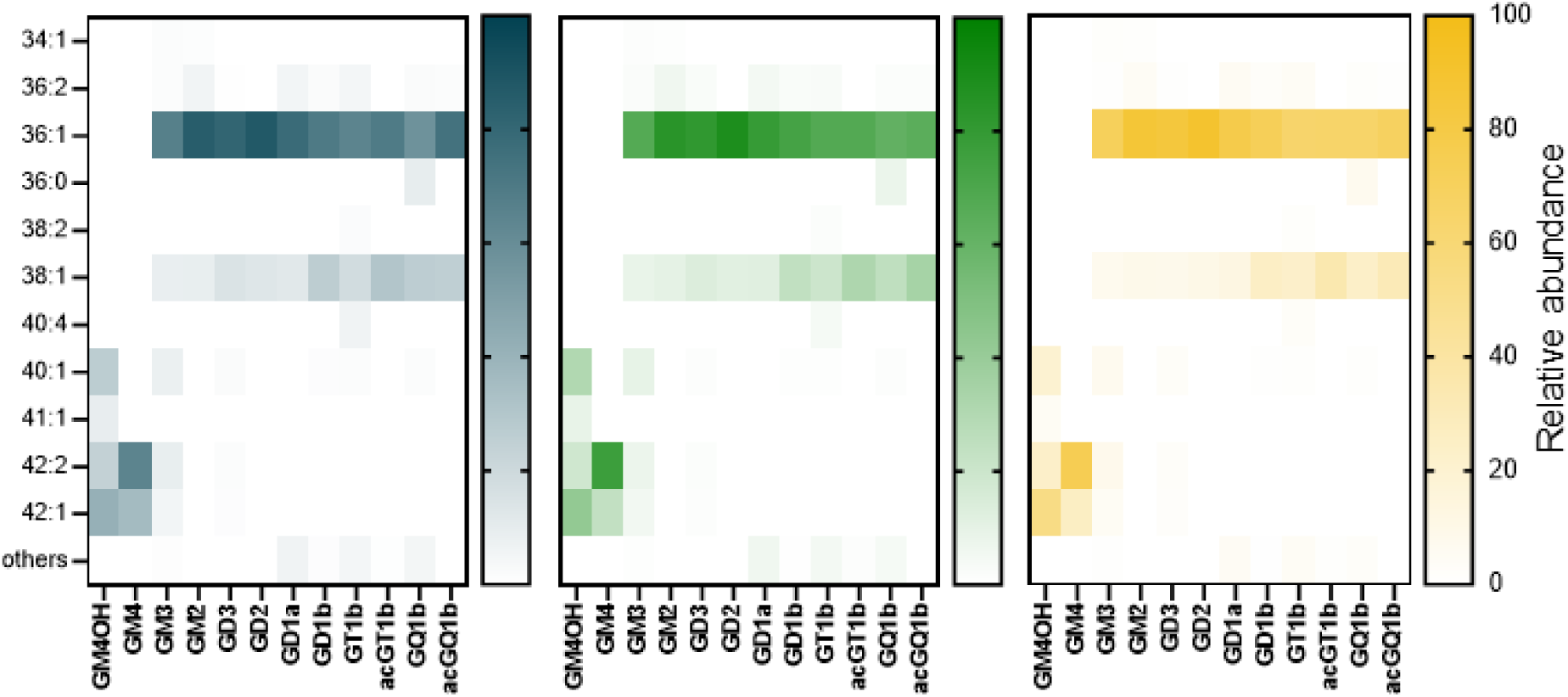
Distribution of the individual species of mouse brain gangliosides. Quantification of the molecular species was performed by liquid chromatography coupled to mass spectrometry (LC-MS) using a triple quadrupole. The QqQ mass spectrometer was operated in negative SRM mode. GG molecular species were identified with theoretical mass and based on previous identification of brain GGs performed in humans using high-resolution LTQ_Orbitrap XL mass spectrometer and MS/MS (19). The ceramides are abbreviated as total number of carbons:number of double bonds. The proportion of each ceramide species was expressed as a percentage of the sum of individual peak areas of all detected molecular species in its specific GG class, every GG class being considered separately. Molecular species which accounted for less than 1% in all three diet conditions were grouped under “others”. Results correspond to the mean of 5-6 mice. The color intensity of a spot on the graph is proportional to the mean percentage of the ceramide species considered within a GG class.

GG molecular species carrying PUFAs were specifically looked for in LC-MS operated in the SRM mode. Some species exhibiting m/z and retention time corresponding to PUFA-carrying GGs were detected as traces in some of the major GG classes of mouse brain: GD1a, GT1b and GQ1b (see Supplementary table 6) but their identity could not be confirmed by MS/MS due to their very low abundance. The diets had no effect on their level.

## 4 Discussion

In order to get insight into the biochemical mechanisms underlying the biological effects of dietary n-3 modulation on the brain, we characterized PL and GG profile under n-3 deficient and n-3 supplemented diets. These two families of complex membrane lipids share some common features and also exhibit major differences. The present study illustrates structural specificities and response to dietary intervention.

Brain PLs, especially PEs, contain high levels of LC-PUFAs. We measured here in mice that C22:6n-3 (DHA) and C20:4n-6 (ARA) levels reach almost 15% and 10% of total brain FAs, respectively, while C16:0, C18:0 and C18:1n-9 still account each for about 20% of total FAs. In PEs, PE 18:0/22:6 is the most abundant molecular species, followed by PE 18:0/20:4. Modulation of dietary n-3 induced significant changes in the distribution of specific FAs among PLs. The n-3 deficient diet, characterized by a high LA/ALA, led to decreased levels of DHA and increased levels of n-6 DPA, as well as the other n-6 family members ARA and C22:4n-6. The n-3 supplemented diet induced the reverse. Impressively, such a subtle change in PL composition, *i.e.* a single double bond difference between DHA and n-6 DPA, has already been shown to have significant consequences on cellular metabolism. Indeed, the requirement for a very specific lipid environment has been known for quite a long time for efficient functioning of visual transduction in the retina through rhodopsin. In the outer segment of photoreceptors, this G protein-coupled receptor relies on the presence of DHA to make a conformational change which is responsible for the induction of visual transduction (37–39). It has been shown that the functional efficacy of rhodopsin was reduced by 50% in model membranes containing n-6 DPA compared with DHA-containing membranes (38). Membranes rich in DHA have unique physical properties, in terms of packing, order and flexibility parameters (39). Specific DHA requirement has also been shown for the brain. LA and n-6 DPA supplementation of rat pups using artificial rearing led to a replacement of brain DHA by n-6 DPA which was associated with a loss in spatial task performance indicating a highly specific structural requirement for DHA (40). In mice, the use of a n-3 deficient diet from gestation to adulthood resulted in increased n-6 DPA levels in brain PLs, at the expense of DHA, which was associated with aberrant wiring of dopaminoreceptive neurons of the striatum, which could account for motivational impairments at adulthood (25). The authors of this study later reported that DHA and n-6 DPA actually interact and modulate the dopamine D2 receptor very differently based on observations using cell membrane extracts and model membranes (26). Indeed, membrane enrichment in DHA, but not n-6 DPA, enhanced ligand binding for various agonists and antagonists. Molecular dynamics simulations indicated that the receptor preferentially interacts with DHA tails suggesting a direct allosteric effect of DHA.

Looking into GGs, we show here in mouse brain that, contrary to PLs, GGs contain mainly saturated FAs with C18:0 accounting for almost 90% of the purified GG FAs detected by GC-FID analysis. LC-MS analyses confirmed this information since ceramide 36:1 was by far the major molecular species detected, corresponding to the combination of the major LCB found in mammals, sphingosine (d18:1), with FA 18:0. d20:1 being the second most present LCB in nervous tissues (36, 41), ceramide 38:1 (d20:1, C18:0) consistently also appeared in our study as an important molecular species in the various GG classes. The predominance of C18:0 is also a feature of SMs, which are, as GGs, ceramide-containing lipids. These observations are in accordance with the ones reported by several other groups studying mouse brain GGs using various LC-MS-based techniques (34, 35, 42). Beside 36:1 and 38:1, minor molecular species are frequently also detected. Here, we measured 34:1, 36:2, 38:2, 40:1, 41:1, 42:2. Those species had already been detected in brain tissues in various proportions and attributed to FAs 16:0, 18:1, 20:1, 22:0, 23:0 and 24:1 thanks to MS/MS characterization (19, 34, 35, 42). GM4 and GM4-OH exhibit a totally different ceramide profile from the other GG classes since they are characterized by the presence of ceramide over 40 carbons carrying one or two unsaturations and hydroxylated fatty acids (C40:1, C42:2, C42:1). This is comparable to the profile described for mouse brain sulfatides (34, 42), which are, as is GM4, derived from galactosylceramide, contrary to the ganglio-series GGs which are derived from glucosylceramide.

Unlike PLs, mouse brain GGs contain no PUFAs. We specifically looked for ceramide species containing PUFAs using MRM mode and detected only traces amounts of GG ceramide species with m/z and retention time corresponding to 38:6, 38:5 and 40:7 and 40:6. No such species had previously been reported in brain to our knowledge, even in studies using high sensitivity MS techniques (43). Here, due to their low levels, characterization of these rare species using MS/MS analyses could not be performed to confirm their identity. GGs are broadly considered to contain saturated FAs and the presence of PUFAs is hardly ever reported. Especially, LA and ALA have been mentioned as minor constituents of the ceramide moiety of GGs in bovine and human milk (44) The virtual absence of PUFAs in GG structure reflects their known role within cellular membranes and the importance of their ceramide moiety for the formation and stabilization of lipid membrane domains. The saturated alkyl chains embedded into the outer leaflet of the plasma membrane makes ceramide a rigid structure reducing membrane fluidity and is one of the GG features which contributes to microdomain structure and function (17). Supporting this idea, Nakagawa *et al.* showed that artificial GM1 carrying ALA, synthesized using a sphingolipid ceramide N-deacylase system, affected the structure of membrane microdomains and induced apoptosis in neuronal cell culture unlike GM1 carrying C18:0 or C18:1 (45).

It appeared in our study that the FA profile of GGs is not modified by LC-PUFA dietary intake, contrary to the FA profile of PLs. The intake of SDA, EPA, DHA through the n-3 supplemented diet did not result in the presence of these FAs in GG structure. Therefore, substrate availability does not seem to be a major determinant of GG FA profile. Specific FA composition of GGs likely results from multiple levels of regulation: substrate preferences of several enzymes including ceramide synthases, glucosyl- and galactosylceramide synthases, sialyltransferases; substrate availability; selective trafficking (specificity of the ceramide transport protein CERT which carries ceramide from the cytosolic side of the endoplasmic reticulum to the trans-Golgi, where it can be glycosylated (24)). However, the precise mechanisms and determinants remain largely unknown. In mammals, ceramides are synthesized by a family of six enzymes called ceramide synthases using FAs usually ranging from 14 to 26 carbons, saturated or mono-unsaturated. The substrate specificity of these enzymes thus initially determines the absence of PUFAs into the structure of glycosphingolipids which are ceramide derivatives. Interestingly, it has been shown that hexosylceramides (glucosyl- and lactosylceramides) which are direct precursors of GGs, contain mainly C22:0, C24:0 and C24:1 in mouse brain. Since brain ceramides mainly carry C18:0 FAs, this suggests that hexosylceramides carrying C18:0 have been selectively used to be further converted into GGs by sialyltransferases with restricted substrate specificity (46).

Overall, our results suggest that the known biological effects of dietary n-3 manipulation on the nervous system does not directly involve brain GGs since their level, profile or ceramide composition are not affected by n-3 deficiency or supplementation. Nevertheless, one cannot exclude an indirect contribution of GGs through changes of distribution within the plasma membrane and disruption of lipid microdomains due to changes in PL structural composition. Indeed, as mentioned earlier, GGs are enriched in membrane microdomains and GG signaling functions are tightly linked to their presence in these domains. Their clustering with signaling proteins such as growth factor receptors favors interaction and regulation of biological processes such as cell proliferation or differentiation (14, 18, 47). Lipid microdomains are thought to play important roles in physiological and pathological events in the nervous system (48). Modulation of microdomain composition and functionality has been proposed as a mechanism through which dietary n-3 PUFAs could exert their beneficial effects (9–11). A line of evidence, obtained *via in vitro* and *in vivo* studies, suggests that LC-PUFAs in membrane PLs affect raft dynamics and consecutively downstream signaling by inducing reorganization of lipid components within the plasma membrane, especially cholesterol and SM. LC-PUFAs cannot pack efficiently with cholesterol and the saturated acyl chains of sphingolipids and PLs that are enriched in microdomains and therefore lead to their displacement thus impacting signaling protein partitioning. It remains unclear whether LC-PUFAs incorporate into and disrupt the clustering of lipid rafts or localize into non raft domains favoring cholesterol displacement into larger and more ordered microdomains. The consequences on GG partitioning and associated signaling is not known either.

In conclusion, n-3 LC-PUFAs dietary intake significantly modifies the PL composition of the mouse brain but do not affect the profile of GGs, which do not contain LC-PUFAs. This suggests that the biological effects of dietary n-3 modulation on neurological processes do not directly involve brain GGs.

## Abbreviations

ALA: α-linolenic acid
ARA: arachidonic acid
DHA: docosahexaenoic
DMA: dimethylacetals
DPA: docosapentaenoic acid
EPA: eicosapentaenoic acid
FA: fatty acid
FAME: fatty acid methyl ester
GC-FID: gas chromatography coupled to a flame ionization detector
GG: ganglioside; GM mono-sialoganglioside
GD: di-sialoganglioside
GT: tri-sialoganglioside
GQ: terta-sialoganglioside
HILIC: hydrophilic interaction liquid chromatography
HPLC: high-performance liquid chromatography
HPTLC: high performance thin layer chromatography
LA: linoleic acid; LC liquid chromatography
LC-MS: liquid chromatography coupled to mass spectrometry
LCB: long chain base
LC-PUFA: Long chain polyunsaturated fatty acid
MS: mass spectrometry
PC: phosphatidylcholine
PE: phosphatidylethanolamine
PL: glycerophospholipid
SA: sialic acid; SDA stearidonic acid
SM: sphingomyelin
SRM: Selected Reaction Monitoring

## Data availability

Data supporting the conclusion of the present study are available from the corresponding author upon reasonable request.

## Ethics Approval

All animal care and experimental procedures were in accordance with the INRA Quality Reference System and to french legislations (Directive 87/148, Ministère de l’Agriculture et de la Pêche) and European (Directive 86/609/EEC). They followed ethical protocols approved by the Region Aquitaine Veterinary Services (Direction Départementale de la Protection des Animaux, approval ID: B33-063-920) and by the animal ethic committee of Bordeaux CEEA50. Every effort was made to minimize suffering and reduce the number of animals used.

## CRediT authorship contribution statement

**E.A.Y.M.**: conceptualization, project administration, funding acquisition, supervision, formal analysis, visualization, writing – original draft; **L.M.**: methodology, investigation, validation, writing – review & editing; **S.C.**: methodology, investigation, validation, data curation, writing - review & editing; **S.G.**: methodology, investigation, validation, writing - review & editing; **F.D.**: methodology, investigation, writing - review & editing; **P.T.**: conceptualization, project administration, funding acquisition, supervision, writing - review & editing; **N.A.**: conceptualization, funding acquisition, writing - review & editing; **O.B.**: methodology, validation, resources, supervision, writing - review & editing.

## Acknowledgments

This work was performed using CALIS network’s resources (doi: 10.15454/9PGN-W156).

