## Supplementary tables for "The ganglioside profile of mouse brain does not respond to changes in dietary n-3 polyunsaturated fatty acid supply"

**Supplementary table 1. Fatty acid composition of the mouse brain (% of total FAs)**

| | Control (balanced) | | n-3 PUFA-deficient | | n-3 PUFA-supplemented | |
|---|---|---|---|---|---|---|
| | **mean** | sd | **mean** | sd | **mean** | sd |
| **C14:0** | **0.13** | *0.01* | **0.12** | *0.01* | **0.15** | *0.01* |
| **DMA16:0** | **2.18** | *0.13* | **2.22** | *0.05* | **2.26** | *0.13* |
| **C16:0** | **20.50** | *0.79* | **20.47** | *0.45* | **20.29** | *0.42* |
| **C16:1n-9** | **0.20** | *0.02* | **0.19** | *0.01* | **0.17** | *0.01* |
| **C16:1n-7** | **0.56** | *0.04* | **0.52** | *0.03* | **0.69** | *0.03* |
| **C17:0** | **0.14** | *0.02* | **0.13** | *0.00* | **0.14** | *0.03* |
| **DMA18:0** | **3.98** | *0.06* | **3.90** | *0.10* | **4.12** | *0.06* |
| **DMA18:1n-9** | **1.41** | *0.06* | **1.28** | *0.02* | **1.36** | *0.06* |
| **DMA18:1n-7** | **1.55** | *0.14* | **1.62** | *0.08* | **1.57** | *0.22* |
| **C18:0** | **19.96** | *0.12* | **20.25** | *0.30* | **19.76** | *0.21* |
| **C18:1t** | **0.17** | *0.03* | **0.11** | *0.03* | **0.13** | *0.03* |
| **C18:1n-9** | **16.05** | *0.27* | **14.82** | *0.09* | **17.57** | *0.43* |
| **C18:1n-7** | **3.61** | *0.15* | **3.62** | *0.08* | **3.57** | *0.09* |
| **C18:2n-6** | **0.32** | *0.01* | **0.46** | *0.03* | **0.30** | *0.01* |
| **C20:0** | **0.29** | *0.05* | **0.26** | *0.03* | **0.26** | *0.03* |
| **C20:1n-9** | **1.36** | *0.13* | **1.22** | *0.04* | **1.32** | *0.11* |
| **C18:3n-3** | **0.33** | *0.02* | **0.31** | *0.02* | **0.34** | *0.02* |
| **C20:2n-6** | **0.07** | *0.01* | **0.10** | *0.01* | **0.05** | *0.01* |
| **C20:3n-9** | **0.23** | *0.02* | **0.11** | *0.01* | **0.24** | *0.02* |
| **C22:0** | **0.20** | *0.03* | **0.19** | *0.02* | **0.21** | *0.02* |
| **C20:3n-6** | **0.33** | *0.05* | **0.25** | *0.01* | **0.44** | *0.02* |
| **C22:1n-9** | **0.18** | *0.01* | **0.16** | *0.00* | **0.17** | *0.02* |
| **C20:4n-6** | **9.44** | *0.23* | **10.51** | *0.29* | **6.48** | *0.25* |
| **C24:0** | **0.32** | *0.04* | **0.25** | *0.03* | **0.25** | *0.03* |
| **C20:5n-3** | **0.00** | *0.00* | **0.00** | *0.00* | **0.51** | *0.03* |
| **C24:1n-9** | **0.44** | *0.05* | **0.45** | *0.05* | **0.50** | *0.09* |
| **C22:4n-6** | **2.29** | *0.06* | **3.03** | *0.10* | **0.98** | *0.04* |
| **C22:5n-6** | **0.66** | *0.17* | **6.03** | *0.18* | **0.12** | *0.01* |
| **C22:5n-3** | **0.12** | *0.01* | **0.05** | *0.00* | **0.63** | *0.05* |
| **C22:6n-3** | **13.02** | *0.07* | **7.38** | *0.28* | **15.44** | *0.65* |
| **Σ SFAs** | **41.54** | 0.73 | **41.67** | 0.38 | **41.06** | 0.49 |
| **Σ MUFAs** | **22.55** | 0.47 | **21.10** | 0.24 | **24.11** | 0.56 |
| **Σ LC-PUFAs** | **26.81** | 0.32 | **28.21** | 0.33 | **25.53** | 0.77 |
| **Σ DMAs** | **9.12** | 0.12 | **9.02** | 0.12 | **9.31** | 0.25 |
| **Σ n-6** | **13.11** | 0.33 | **20.37** | 0.41 | **8.37** | 0.24 |
| **Σ n-3** | **13.47** | 0.08 | **7.73** | 0.27 | **16.91** | 0.69 |
| **n-6/n-3** | **0.97** | 0.03 | **2.63** | 0.13 | **0.50** | 0.02 |

Brain total lipids of mice subjected to the different diets were analyzed using GC-FID after trans-methylation. Results correspond to the relative level of each detected FA expressed as a percentage of total FAs (FA methyl esters (FAMEs) and dimethylacetals (DMAs)) and are given as mean ± SD of 5-6 mice. SFAs, saturated fatty acids; MUFAs, monounsaturated fatty acids; LC-PUFAs, long chain polyunsaturated fatty acids; DMAs, dimethylacetals

**Supplementary table 2. Distribution of the individual species of mouse brain phosphatidylethanolamines (% of total PEs)**

| | | | Control (balanced) | | n-3 PUFA-deficicent | | n-3 PUFA-supplemented | |
|---|---|---|---|---|---|---|---|---|
| | | $[M+H]^+$ | **mean** | sd | **mean** | sd | **mean** | sd |
| **PE 34:2** | **PE 16:1/18:1**<br>***PE 16:0/18:2*** | 716.5225 | tr | | tr | | tr | |
| **PE 34:1** | **PE 16:0/18:1** | 718.5381 | **5.0** | 0.52 | **4.6** | 0.39 | **5.6** | 0.38 |
| **PE 34:0** | **PE 16:0/18:0** | 720.5538 | tr | | tr | | tr | |
| **PE 36:5** | **PE 16:1/20:4** | 738.5068 | tr | | tr | | tr | |
| **PE 36:4** | **PE 16:0/20:4** | 740.5225 | **2.4** | 0.40 | **2.7** | 0.26 | **1.4** | 0.09 |
| **PE 36:3** | **PE 18:1/18:2**<br>***PE 16:0/20:3*** | 742.5381 | tr | | tr | | tr | |
| **PE 36:2** | **PE 18:1/18:1** | 744.5538 | **6.8** | 1.65 | **5.4** | 0.11 | **8.7** | 0.77 |
| **PE 36:1** | **PE 18:0/18:1** | 746.5694 | **7.8** | 0.90 | **7.5** | 0.33 | **7.3** | 0.73 |
| **PE 38:7** | **PE 16:1/22:6** | 762.5068 | tr | | tr | | tr | |
| **PE 38:6** | **PE 16:0/22:6** | 764.5225 | **9.2** | 1.69 | **5.0** | 0.23 | **10.9** | 0.38 |
| **PE 38:5** | **PE 18:1/20:4**<br>***PE 16:0/22:5*** | 766.5381 | **3.2** | 0.46 | **7.5** | 0.47 | **2.3** | 0.19 |
| **PE 38:4** | **PE 18:0/20:4**<br>***PE 16:0/22:4 tr*** | 768.5538 | **18.7** | 2.46 | **21.3** | 1.09 | **12.7** | 0.40 |
| **PE 38:3** | **PE 18:0/20:3** | 770.5694 | tr | | tr | | tr | |
| **PE 38:2** | **PE 18:1/20:1** | 772.5851 | **0.6** | 0.16 | **0.5** | 0.09 | **0.5** | 0.17 |
| **PE P-40:7** | **PE P-18:1/22:6** | 774.5432 | **0.9** | 0.22 | **0.9** | 0.07 | **0.8** | 0.13 |
| **PE 38:0**<br>**PE P-40:6** | **PE 18:0/20:0**<br>**PE P-18:0/22:6** | 776.6164<br>776.5589 | **0.9** | 0.15 | **0.5** | 0.10 | **1.1** | 0.14 |
| **PE 40:9** | NI | 786.5068 | tr | | tr | | tr | |
| **PE 40:8** | NI | 788.5225 | tr | | tr | | tr | |
| **PE 40:7** | **PE 18:1/22:6** | 790.5381 | **3.1** | 0.47 | **1.7** | 0.17 | **3.3** | 0.16 |
| **PE 40:6** | **PE 18:0/22:6** | 792.5539 | **33.3** | 2.88 | **19.7** | 1.00 | **39.4** | 1.81 |
| **PE 40:5** | **PE 18:0/22:5** | 794.5694 | **1.8** | 1.03 | **15.1** | 0.52 | **0.4** | 0.56 |
| **PE 40:4** | **PE 18:0/22:4** | 796.5851 | **3.5** | 0.39 | **4.4** | 0.48 | **1.3** | 0.20 |
| **PE 42:9** | **PE 20:4/22:5** | 814.5381 | tr | | tr | | tr | |
| **PE 42:8** | **PE 20:4/22:4** | 816.5538 | tr | | tr | | tr | |
| **PE 42:7** | NI | 818.5694 | tr | | tr | | tr | |
| **PE 44:12** | **PE 22:6/22:6** | 836.5225 | tr | | tr | | tr | |
| **PE 44:10** | **PE 22:4/22:6** | 840.5538 | tr | | tr | | tr | |

Brain PEs of mice subjected to the different diets were analyzed using liquid chromatography coupled to mass spectrometry (LC-MS). Quantification of the molecular species was performed using a triple quadrupole and identification using a Orbitrap Fusion. Species are abbreviated as total number of carbons:number of double bonds and when identified, position on the glycerol backbone is shown as sn-1/sn-2 of the fatty acid radicals. Theoretical exact molecular mass of the protonated molecular ion ($[M+H]^+$) is given. Minor isobaric species are written in *italics*. PE 38:0 and PE P-40:6 could not be distinguished with the triple quadrupole spectrometer due to insufficient resolution. Results correspond to the relative level of each detected molecular species expressed as a percentage of the sum of all molecular species signal intensities. Results are given as mean ± SD of 5-6 mice. PE P-, phosphatidylethanolamine vinyl-ether (plasmalogen); tr, traces; NI, non-identified using Orbitrap Fusion.

**Supplementary table 3. Distribution of the individual species of mouse brain phosphatidylcholines (% of total PCs)**

| | | | Control (balanced) | | n-3 PUFA-deficicent | | n-3 PUFA-supplemented | |
|---|---|---|---|---|---|---|---|---|
| | | [M+H]+ | **mean** | sd | **mean** | sd | **mean** | sd |
| **PC 30:0** | **PC 16:0/14:0** | 706.5381 | tr | | tr | | tr | |
| **PC P-32:0** | **PC P-16:0/16:0**<br>**PC O-16:1/16:0** | 718.5745 | tr | | tr | | tr | |
| **PC 31:0**<br>**PC O-32:0** | **PC 15:0/16:0**<br>**PC O-16:0/16:0** | 720.5538<br>720.5902 | tr | | tr | | tr | |
| **PC 32:2** | NI | 730.5381 | tr | | tr | | tr | |
| **PC 32:1** | **PC 16:0/16:1**<br>***PC 14:0/18:1*** | 732.5538 | **1.7** | 0.23 | **1.5** | 0.12 | **1.6** | 0.14 |
| **PC 32:0** | **PC 16:0/16:0** | 734.5694 | **23.0** | 3.01 | **21.4** | 0.95 | **20.3** | 0.32 |
| **PC P-34:1**<br>**PC O-34:0** | NI | 744.5902 | tr | | tr | | tr | |
| **PC O-34:1** | **PC O-16:0/18:1**<br>**PC O-18:0/16:1** | 746.6058 | **0.7** | 0.18 | **0.5** | 0.09 | **0.6** | 0.08 |
| **PC 34:2** | **PC 16:1/18:1**<br>***PC 16:0/18:2*** | 758.5694 | **0.6** | 0.11 | **0.7** | 0.08 | **0.6** | 0.07 |
| **PC 34:1** | **PC 16:0/18:1** | 760.5851 | **32.2** | 2.36 | **30.6** | 0.55 | **32.5** | 0.32 |
| **PC 34:0** | **PC 16:0/18:0** | 762.6007 | **4.8** | 0.63 | **5.0** | 0.28 | **5.1** | 0.26 |
| **PC P-36:2** | NI | 770.6058 | tr | | tr | | tr | |
| **PC P-36:1** | **PC P-18:0/18:1** | 772.6215 | tr | | tr | | tr | |
| **PC P-36:0** | **PC P-18:0/18:0** | 774.6371 | tr | | tr | | tr | |
| **PC 36:4** | **PC 16:0/20:4** | 810.6007 | **4.1** | 0.86 | **4.9** | 0.28 | **2.6** | 0.22 |
| **PC 36:3** | **PC 16:0/20:3** | 784.5851 | tr | | tr | | tr | |
| **PC 36:2** | **PC 18:1/18:1** | 786.6007 | **2.9** | 0.34 | **2.7** | 0.15 | **3.0** | 0.20 |
| **PC 36:1** | **PC 18:0/18:1**<br>***PC 16:0/20:1 tr*** | 788.6164 | **11.8** | 1.36 | **12.2** | 0.50 | **12.9** | 0.21 |
| **PC P-38:6** | NI | 790.5745 | tr | | tr | | tr | |
| **PC P-38:0** | NI | 802.6684 | tr | | tr | | tr | |
| **PC 38:7** | **PC 16:1/22:6** | 804.5538 | tr | | tr | | tr | |
| **PC 38:6** | **PC 16:0/22:6** | 806.5694 | **3.5** | 0.51 | **1.8** | 0.05 | **5.3** | 0.40 |
| **PC 38:5** | **PC 18:1/20:4** | 808.5851 | **1.6** | 0.19 | **3.5** | 0.15 | **1.2** | 0.06 |
| **PC 38:4** | **PC 18:0/20:4** | 810.6007 | **4.9** | 0.40 | **5.9** | 0.12 | **3.4** | 0.21 |
| **PC 38:3** | **PC 18:0/20:3** | 812.6164 | tr | | tr | | tr | |
| **PC 38:2** | **PC 18:1/20:1** | 814.6320 | tr | | tr | | tr | |
| **PC 38:1** | **PC 18:1/20:0**<br>**PC 18:0/20:1** | 815.6399 | tr | | tr | | tr | |
| **PC P-40:6** | **PC P-18:0/22:6** | 818.6058 | tr | | tr | | tr | |
| **PC 40:7** | **PC 18:1/22:6** | 832.5851 | **0.7** | 0.19 | **0.4** | 0.05 | **1.0** | 0.09 |
| **PC 40:6** | **PC 18:0/22:6** | 834.6007 | **1.8** | 0.45 | **1.5** | 0.14 | **2.6** | 0.25 |
| **PC 40:5** | **PC 18:0/22:5**<br>**PC 20:1/20:4**<br>***PC 18:1/22:4*** | 836.6164 | tr | | tr | | tr | |
| **PC 40:4** | **PC 18:0/22:4** | 838.6320 | tr | | tr | | tr | |

Brain PCs of mice subjected to the different diets were analyzed using liquid chromatography coupled to mass spectrometry (LC-MS). Quantification of the molecular species was performed using a triple quadrupole and identification using a Orbitrap Fusion. Species are abbreviated as total number of carbons:number of double bonds and when identified, position on the glycerol backbone is shown as sn-1/sn-2 of the fatty acid radicals. Theoretical exact molecular mass of the protonated molecular ion ($[M+H]^+$) is given. Minor isobaric species are written in *italics*. Results correspond to the relative level of each detected molecular species expressed as a percentage of the sum of all molecular species signal intensities. Results are given as mean ± SD of 5-6 mice. PC P-, phosphatidylcholine vinyl-ether (plasmalogen); PC O-, phosphatidylcholine alkyl-ether; tr, traces; NI, non-identified using Orbitrap Fusion.

**Supplementary table 4. Distribution of the individual species of mouse brain sphingomyelins (% of total SMs)**

| | | | Control (balanced) | | n-3 PUFA-deficient | | n-3 PUFA-supplemented | |
|---|---|---|---|---|---|---|---|---|
| | | $[M+H]^+$ | **mean** | sd | **mean** | sd | **mean** | sd |
| **SM 32:1** | NI | 675.5436 | tr | | tr | | tr | |
| **SM 33:1** | NI | 689.5592 | tr | | tr | | tr | |
| **SM 34:2** | NI | 701.5592 | tr | | tr | | tr | |
| **SM 34:1** | **d18:1/16:0** | 703.5749 | **2.7** | 1.24 | **2.0** | 0.40 | **2.2** | 0.46 |
| **SM 35:1** | **d18:1/17:0** | 717.5905 | tr | | tr | | tr | |
| **SM 36:4** | NI | 725.5592 | tr | | tr | | tr | |
| **SM 36:3** | NI | 727.5749 | tr | | tr | | tr | |
| **SM 36:2** | **d18:1/18:1** | 729.5905 | **3.9** | 1.04 | **5.4** | 0.50 | **4.9** | 1.17 |
| **SM 36:1** | **d18:1/18:0** | 731.6062 | **70.8** | 4.38 | **67.8** | 2.08 | **67.8** | 3.25 |
| **SM 37:1** | **d18:1/19:0** | 745.6218 | tr | | tr | | tr | |
| **SM 38:2** | **d18:1/20:1** | 757.6218 | tr | | tr | | tr | |
| **SM 38:1** | **d18:1/20:0** | 759.6375 | **4.7** | 1.43 | **5.2** | 0.66 | **5.1** | 0.66 |
| **SM 39:1** | **d18:1/21:0** | 773.6531 | tr | | tr | | tr | |
| **SM 40:2** | **d18:1/22:1** | 785.6531 | **0.5** | 0.42 | **0.9** | 0.21 | **0.8** | 0.33 |
| **SM 40:1** | **d18:1/22:0** | 787.6688 | **2.3** | 0.42 | **1.7** | 0.59 | **1.9** | 0.37 |
| **SM 41:2** | **d18:2/23:0** | 799.6688 | tr | | tr | | tr | |
| **SM 41:1** | **d18:1/23:0** | 801.6844 | **0.6** | 0.47 | **1.0** | 0.27 | **0.8** | 0.37 |
| **SM 42:3** | **d18:2/24:1** | 811.6688 | tr | | tr | | tr | |
| **SM 42:2** | **d18:1/24:1** | 813.6844 | **11.1** | 2.04 | **11.5** | 1.07 | **12.4** | 1.80 |
| **SM 42:1** | **d18:1/24:0** | 815.7001 | **1.6** | 1.09 | **2.6** | 0.75 | **1.9** | 0.67 |
| **SM 44:2** | NI | 841.7157 | tr | | tr | | tr | |

Brain SMs of mice subjected to the different diets were analyzed using liquid chromatography coupled to mass spectrometry (LC-MS). Quantification of the molecular species was performed using a triple quadrupole and identification using a Orbitrap Fusion. The ceramide backbone is abbreviated as total number of carbons:number of double bonds and when identified, the position on the ceramide backbone is shown as long chain base/fatty acid radical. Theoretical exact molecular mass of the protonated molecular ion ($[M+H]^+$) is given. Results correspond to the relative levels of each detected molecular species expressed as a percentage of the sum of all molecular species signal intensities and are given as mean ± SD of 5-6 mice. tr, traces; NI, non-identified using Orbitrap Fusion.

**Supplementary table 5. Fatty acid composition of mouse brain gangliosides (% of total FAs)**

| | **Control (balanced)** | | **n-3 PUFA-deficient** | | **n-3 PUFA-supplemented** | |
|---|---|---|---|---|---|---|
| | **mean** | sd | **mean** | sd | **mean** | sd |
| 16:0 | **3.2** | 1.67 | **2.3** | 0.57 | **2.7** | 2.03 |
| 16:1n-9 | **0.4** | 0.40 | **0.1** | 0.06 | **0.2** | 0.22 |
| 17:0 | **1.1** | 0.25 | **1.1** | 0.27 | **1.1** | 0.63 |
| 18:0 | **86.9** | 2.61 | **88.0** | 0.95 | **86.5** | 3.25 |
| 18:1n-9 | **0.8** | 0.28 | **1.7** | 0.74 | **1.3** | 1.19 |
| 18:2n-6 | **0.6** | 0.99 | **0.3** | 0.04 | **0.7** | 1.19 |
| 20:0 | **5.6** | 0.45 | **5.1** | 0.32 | **6.0** | 1.15 |
| 22:0 | **1.0** | 0.28 | **0.9** | 0.19 | **1.0** | 0.23 |
| 24:0 | **0.5** | 0.14 | **0.5** | 0.30 | **0.5** | 0.12 |

After hydrolysis of phospholipids and extraction of gangliosides from total lipid extracts, fatty acids of the brain gangliosides of mice subjected to the different diets were analyzed using GC-FID (gas chromatography coupled to a flame ionization detector) after trans-methylation. Results correspond to the relative level of each detected fatty acid expressed as a percentage of the sum of all fatty acids signal intensities and are given as mean ± SD of 5-6 mice.

**Supplementary table 6. Distribution of the individual species of mouse brain gangliosides**

| | | Control (balanced) | | n-3 PUFA-deficicent | | n-3 PUFA-supplemented | |
|---|---|---|---|---|---|---|---|
| | | **mean** | sd | **mean** | sd | **mean** | sd |
| **GM4OH** | [M-H]$^{-}$ | | | | | | |
| 40:1 | 1089.7419 | **26.2** | 13.98 | **30.3** | 6.93 | **19.6** | 11.76 |
| 41:1 | 1103.7575 | **9.0** | 7.90 | **8.8** | 5.38 | **4.5** | 2.96 |
| 42:2 | 1115.7575 | **23.7** | 10.68 | **18.6** | 4.83 | **23.2** | 14.07 |
| 42:1 | 1117.7732 | **41.2** | 16.31 | **42.3** | 10.68 | **52.7** | 14.05 |
| **GM4** | [M-H]$^{-}$ | | | | | | |
| 42:2 | 1099.7626 | **64.4** | 13.40 | **76.1** | 4.66 | **74.3** | 7.40 |
| 42:1 | 1101.7783 | **35.6** | 13.40 | **23.9** | 4.66 | **25.7** | 7.40 |
| **GM3** | [M-H]$^{-}$ | | | | | | |
| 34:1 | 1151.7059 | **1.3** | 0.67 | **0.9** | 0.49 | **1.5** | 0.86 |
| 36:2 | 1177.7215 | **1.2** | 0.80 | **1.9** | 1.35 | **0.7** | 0.51 |
| 36:1 | 1179.7372 | **66.7** | 5.28 | **66.9** | 3.52 | **70.8** | 3.38 |
| 38:1 | 1207.7685 | **8.5** | 0.91 | **8.3** | 2.31 | **7.3** | 2.65 |
| 40:2 | 1233.7841 | tr | | tr | | tr | |
| 40:1 | 1235.7998 | **7.8** | 2.47 | **9.4** | 4.22 | **6.8** | 2.76 |
| 42:2 | 1261.8154 | **8.9** | 2.16 | **7.2** | 2.72 | **7.8** | 2.61 |
| 42:1 | 1263.8311 | **4.9** | 1.15 | **4.9** | 2.51 | **4.5** | 1.19 |
| **GM2** | [M-H]$^{-}$ | | | | | | |
| 34:1 | 1354.7853 | **1.0** | 0.79 | **0.6** | 0.36 | **1.0** | 1.09 |
| 36:2 | 1380.8009 | **5.4** | 3.55 | **5.6** | 2.19 | **3.9** | 2.11 |
| 36:1 | 1382.8166 | **84.8** | 5.11 | **83.7** | 3.23 | **86.3** | 4.22 |
| 38:1 | 1410.8479 | **8.8** | 1.79 | **10.1** | 1.77 | **8.8** | 2.93 |
| **GD3** | [M-2H]$^{2-}$ | | | | | | |
| 36:2 | 733.9048 | **0.3** | 0.16 | **2.3** | 1.35 | **0.7** | 0.91 |
| 36:1 | 734.9127 | **81.0** | 8.37 | **80.9** | 8.69 | **83.6** | 3.55 |
| 38:1 | 748.9283 | **14.4** | 5.87 | **13.4** | 9.53 | **8.5** | 4.60 |
| 40:1 | 762.9440 | **1.9** | 1.65 | **1.0** | 1.17 | **2.9** | 2.15 |
| 42:2 | 775.9518 | **1.3** | 0.50 | **1.4** | 0.84 | **2.4** | 1.58 |
| 42:1 | 776.9596 | **1.1** | 0.88 | **0.9** | 0.94 | **1.8** | 1.94 |
| **GD2** | [M-2H]$^{2-}$ | | | | | | |
| 36:1 | 836.4523 | **87.3** | 7.79 | **88.4** | 10.33 | **90.0** | 3.87 |
| 38:1 | 850.4680 | **12.7** | 7.79 | **11.6** | 10.33 | **10.0** | 3.87 |
| **GD1a** | [M-2H]$^{2-}$ | | | | | | |
| 34:1 | 903.4631 | **0.6** | 0.12 | **0.6** | 0.04 | **0.6** | 0.22 |
| 36:2 | 916.4709 | **4.8** | 1.03 | **4.7** | 0.50 | **4.9** | 0.94 |
| 36:1 | 917.4788 | **77.5** | 1.39 | **78.5** | 0.80 | **77.6** | 3.12 |
| 38:6 | 926.4546 | tr | | tr | | tr | |
| 38:2 | 930.4866 | **0.8** | 0.22 | **0.7** | 0.07 | **0.7** | 0.29 |
| 38:1 | 931.4944 | **11.8** | 0.70 | **11.9** | 0.60 | **12.5** | 2.49 |
| 39:1 | 938.5022 | tr | | tr | | tr | |
| 40:7 | 939.4622 | tr | | tr | | tr | |
| 40:6 | 940.4702 | tr | | tr | | tr | |
| 40:5 | 941.5000 | tr | | tr | | tr | |

| | | | | | | | |
|---|---|---|---|---|---|---|---|
| 40:4 | 942.4862 | tr | | tr | | tr | |
| 40:2 | 944.5022 | tr | | tr | | tr | |
| 40:1 | 945.5101 | **0.8** | 0.22 | **0.8** | 0.10 | **0.7** | 0.33 |
| 41:1 | 952.5179 | tr | | tr | | tr | |
| 42:2 | 958.5179 | tr | | tr | | tr | |
| 42:1 | 959.5257 | tr | | tr | | tr | |
| **GD1b** | $[M-2H]^{2-}$ | | | | | | |
| 36:2 | 916.4709 | **1.9** | 0.78 | **2.0** | 0.35 | **2.3** | 1.19 |
| 36:1 | 917.4888 | **69.9** | 3.58 | **72.8** | 2.20 | **72.2** | 3.68 |
| 38:1 | 931.4944 | **25.9** | 2.83 | **24.3** | 2.13 | **24.7** | 3.07 |
| 40:1 | 945.5101 | **1.1** | 0.52 | **0.5** | 0.27 | **0.5** | 0.57 |
| **GT1b** | $[M-2H]^{2-}$ | | | | | | |
| 36:2 | 1062.0188 | **4.4** | 1.93 | **2.4** | 1.75 | **4.0** | 0.50 |
| 36:1 | 1063.0265 | **64.4** | 4.74 | **67.2** | 5.68 | **63.9** | 1.21 |
| 38:6 | 1072.0023 | **0.9** | 0.60 | **0.7** | 0.24 | **1.0** | 0.31 |
| 38:5 | 1073.0103 | **1.0** | 0.64 | **1.3** | 0.84 | **0.7** | 0.39 |
| 38:2 | 1076.0343 | **1.7** | 1.12 | **1.4** | 0.19 | **1.4** | 0.57 |
| 38:1 | 1077.0421 | **18.0** | 4.46 | **19.8** | 3.80 | **22.9** | 0.86 |
| 39:1 | 1084.0499 | tr | | tr | | tr | |
| 40:7 | 1085.0999 | tr | | tr | | tr | |
| 40:6 | 1086.0179 | tr | | tr | | tr | |
| 40:4 | 1088.0339 | **5.8** | 2.73 | **3.8** | 2.39 | **2.9** | 0.82 |
| 40:2 | 1090.0499 | tr | | tr | | tr | |
| 40:1 | 1091.0578 | **1.3** | 0.84 | **0.7** | 0.59 | **1.1** | 0.40 |
| 42:2 | 1104.0656 | tr | | tr | | tr | |
| **GT1b ac** | $[M-2H]^{2-}$ | | | | | | |
| 36:1 | 1084.0317 | **69.2** | 6.85 | **67.4** | 15.48 | **64.6** | 5.33 |
| 38:2 | 1097.0395 | **0.5** | 0.85 | **1.0** | 1.29 | **0.9** | 0.57 |
| 38:1 | 1098.0474 | **30.3** | 6.83 | **31.6** | 14.51 | **34.5** | 5.28 |
| **GQ1b** | $[M-3H]^{3-}$ | | | | | | |
| 36:2 | 804.7085 | **1.9** | 1.01 | **1.3** | 0.49 | **1.4** | 0.78 |
| 36:1 | 805.3804 | **57.3** | 2.46 | **61.5** | 2.65 | **64.7** | 7.22 |
| 36:0 | 806.0522 | **9.2** | 0.87 | **7.6** | 3.05 | **6.7** | 3.03 |
| 38:6 | 811.3642 | tr | | tr | | tr | |
| 38:2 | 814.0522 | tr | | tr | | tr | |
| 38:1 | 814.7241 | **25.8** | 1.99 | **24.9** | 3.15 | **23.8** | 5.10 |
| 39:1 | 819.3960 | tr | | tr | | tr | |
| 40:7 | 820.0360 | tr | | tr | | tr | |
| 40:6 | 820.7080 | tr | | tr | | tr | |
| 40:1 | 824.0679 | **1.2** | 0.77 | **1.0** | 0.53 | **1.1** | 0.59 |
| 41:1 | 828.7398 | tr | | tr | | tr | |
| 42:2 | 832.7398 | tr | | tr | | tr | |
| 42:1 | 833.4117 | tr | | tr | | tr | |
| **GQ1b ac** | $[M-3H]^{3-}$ | | | | | | |
| 36:2 | 818.7122 | **1.4** | 1.06 | **1.4** | 1.09 | **0.5** | 0.79 |
| 36:1 | 819.3840 | **73.7** | 5.64 | **64.1** | 3.44 | **68.7** | 8.85 |
| 38:1 | 828.7278 | **25.0** | 6.10 | **34.6** | 3.82 | **30.8** | 9.04 |

Quantification of the molecular species was performed by liquid chromatography coupled to mass spectrometry (LC-MS) using a triple quadrupole. The QqQ mass spectrometer was operated in negative SRM mode. GG molecular species were identified with theoretical mass and based on previous identification of brain GGs performed in humans using high-resolution LTQ_Orbitrap XL mass spectrometer and MS/MS {Sibille, 2016 #1110}. The ceramides are abbreviated as total number of carbons:number of double bonds. Theoretical exact molecular mass of the deprotonated ion ($[M-xH]^{x-}$) is given. Results correspond to the proportion of each ceramide species expressed as a percentage of the sum of individual peak areas of all detected molecular species in its specific GG class, every GG class being considered separately. Results correspond to the mean of 5-6 mice. tr, traces
